# Putatively cancer-specific alternative splicing is shared across patients and present in developmental and other non-cancer cells

**DOI:** 10.1101/754044

**Authors:** Julianne K. David, Sean K. Maden, Benjamin R. Weeder, Reid F. Thompson, Abhinav Nellore

## Abstract

We compared cancer and non-cancer RNA sequencing (RNA-seq) data from The Cancer Genome Atlas (TCGA), the Genotype-Tissue Expression (GTEx) Project, and the Sequence Read Archive (SRA). We found that: 1) averaging across cancer types, 80.6% of exon-exon junctions thought to be cancer-specific based on comparison with tissue-matched samples are in fact present in other adult non-cancer tissues throughout the body; 2) 30.8% of junctions not present in any GTEx or TCGA normal tissues are shared by multiple samples within at least one cancer type cohort, and 87.4% of these distinguish between different cancer types; and 3) many of these junctions not found in GTEx or TCGA normal tissues (15.4% on average) are also found in embryological and other developmentally associated cells. This study probes the distribution of putatively cancer-specific junctions across a broad set of publicly available non-cancer human RNA-seq datasets. Overall, we identify a subset of shared cancer-specific junctions that could represent novel sources of cancer neoantigens. We further describe a framework for characterizing possible origins of these junctions, including potential developmental and embryological sources, as well as cell type-specific markers particularly related to cell types of cancer origin. These findings refine the meaning of RNA splicing event novelty, particularly with respect to the human neoepitope repertoire. Ultimately, cancer-specific exon-exon junctions may affect the anti-cancer immune response and may have a substantial causal relationship with the biology of disease.

## INTRODUCTION

Aberrant RNA splicing is increasingly recognized as a feature of malignancy (Climente-González et al., 2017; Sebestyén et al., 2015; Srebrow and Kornblihtt, 2006; Sveen et al., 2016; Xiong et al., 2015), with potential prognostic significance across many cancer types including non-small cell lung cancer, ovarian cancer, breast cancer, colorectal cancer, uveal melanoma, and glioblastoma (Bjørklund et al., 2017; Li et al., 2017; Marcelino Meliso et al., 2017; Robertson et al., 2018; Zhu et al., 2018; Zong et al., 2018). Due to its potential for generating novel peptide sequences, aberrant RNA splicing is also interesting as a potential source of neoantigens for cancer immunotherapy targeting. For instance, retained intronic sequences can give rise to numerous potential antigens among patients with melanoma, although they are not a significant predictor of cancer immunotherapy response (Smart et al., 2018), and a patient-specific neoantigen arising from a gene fusion has been shown to lead to complete response from immune checkpoint blockade (Yang et al., 2019). Novel cancer-specific exon-exon junctions have also been shown to be a source of peptide antigens (Kahles et al., 2018), and represent compelling potential targets for personalized anti-cancer vaccines (Slansky and Spellman, 2019).

However, the ability of the adaptive immune system to target a given antigen as “foreign” depends on a complex prior tolerogenic education, and in particular on whether or not a given antigen has been previously “seen” by the immune system in a healthy context (Klein et al., 2014). Therefore, prediction of cancer-specific antigens depends explicitly on their sequence novelty, and thus requires a comparison with non-cancer cells.

Choosing a “normal” tissue standard for comparison is difficult in the context of RNA sequencing (RNA-seq) data analysis, given the presence of alternative splicing throughout normal and cancerous biological processes (Norris and Calarco, 2012; Sveen et al., 2016; Yang et al., 2016). Previously, cancer-specific aberrant splicing has been detected by comparing tumor RNA-seq data against a single reference annotation (Tang and Madhavan, 2017) or a limited “panel of normals” (Smart et al., 2018). More recently, a TCGA network paper (Kahles et al., 2018) used the large publicly available datasets of The Cancer Genome Atlas (TCGA) (Cancer Genome Atlas Research Network et al., 2013) and the Genotype Tissue Expression project (GTEx) (Cancer Genome Atlas Research Network et al., 2013; GTEx Consortium, 2015) to identify and validate thousands of novel splicing events including exon-exon junctions present in a specific TCGA cancer type but not in the corresponding normal adult tissue in GTEx. This study also predicted alternative splicing neoepitopes (ASNs) via this comparison, and validated several ASNs shared between multiple patients with the intracellular proteomics data available for select ovarian and breast cancer TCGA donors in the Clinical Proteomic Tumor Analysis Consortium dataset (Kahles et al., 2018).

## RESULTS

### Cancers harbor many novel shared exon-exon junctions not present in adult non-cancer tissues or cells

While cancer-specific exon-exon junctions identified using tissue-matched normal samples have the potential to give rise to neoantigens (Kahles et al., 2018), we reasoned that they could be expressed in other normal tissues due to variability in patterns of transcription and alternative splicing among different tissues (Saha et al., 2017). In such cases, these junctions might not yield bona fide neoantigens due to the prior tolerogenic education of the immune system. We therefore re-evaluated the incidence of cancer-specific junctions using RNA-seq data from TCGA and the large compendium of adult tissues from GTEx. We found that on average, across cancer types, 80.6% of junctions thought to be cancer-specific based on comparison only with tissue-matched samples are in fact present in other adult non-cancer tissues and cell types throughout the body. Across cancer types, an average of 90.2% of all junctions found in cancer samples are also present in one or more adult normal samples from GTEx or TCGA [“core normals”] (Figure 1A). The overall number of these novel junctions varies both within and across different cancer types, with ovarian carcinoma and uveal melanoma having the highest and lowest average number of junctions per sample, respectively (Figure 1B, Table S1), although we find that the set of junctions defined as “novel” is highly sensitive to the filtering criteria used (see Figure S1A, Table S2, and “Selection of cancer-specific junctions” in Methods). We use lack of occurrence of a junction in core normals as a baseline definition of cancer specificity.

**Figure 1:**
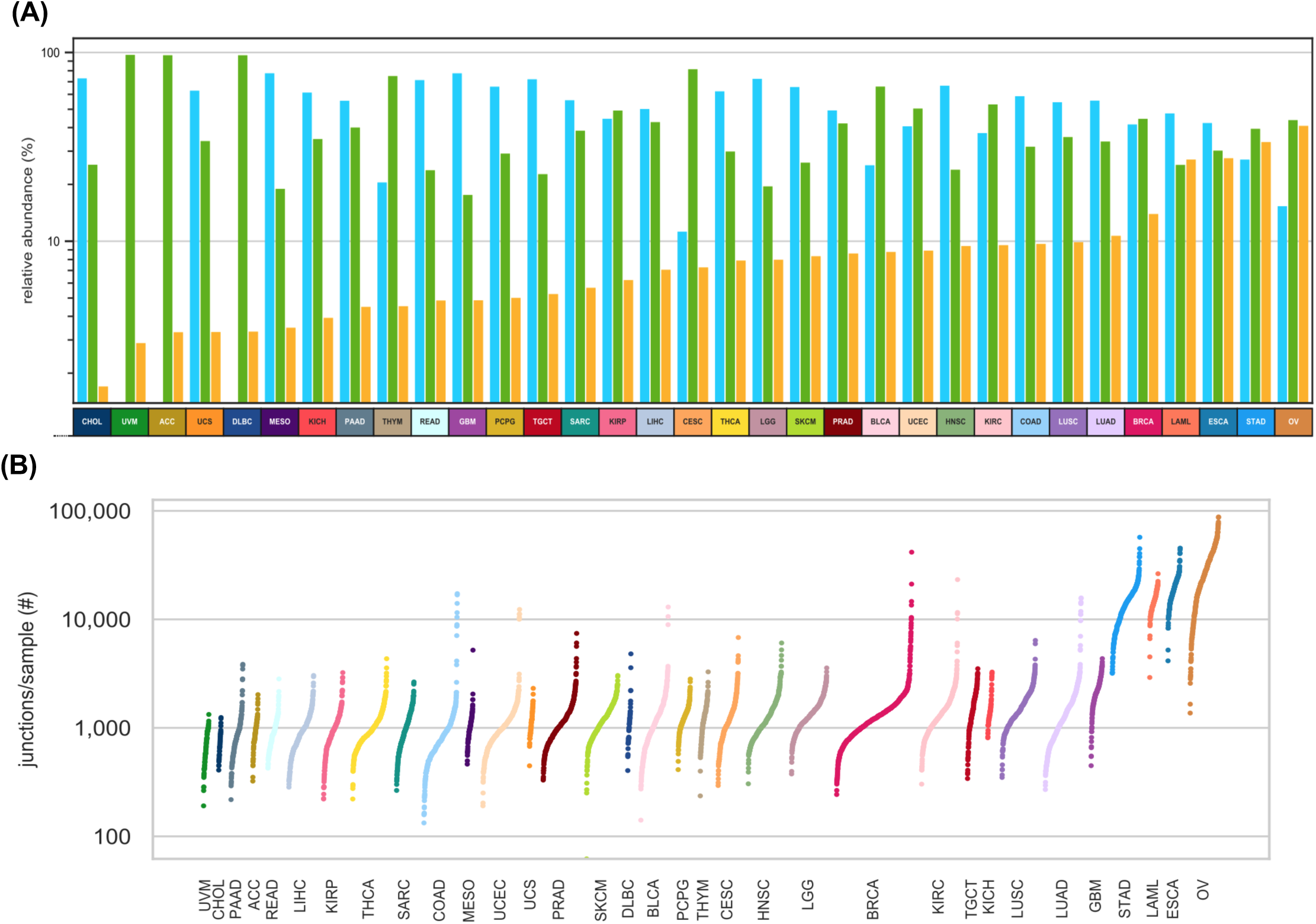
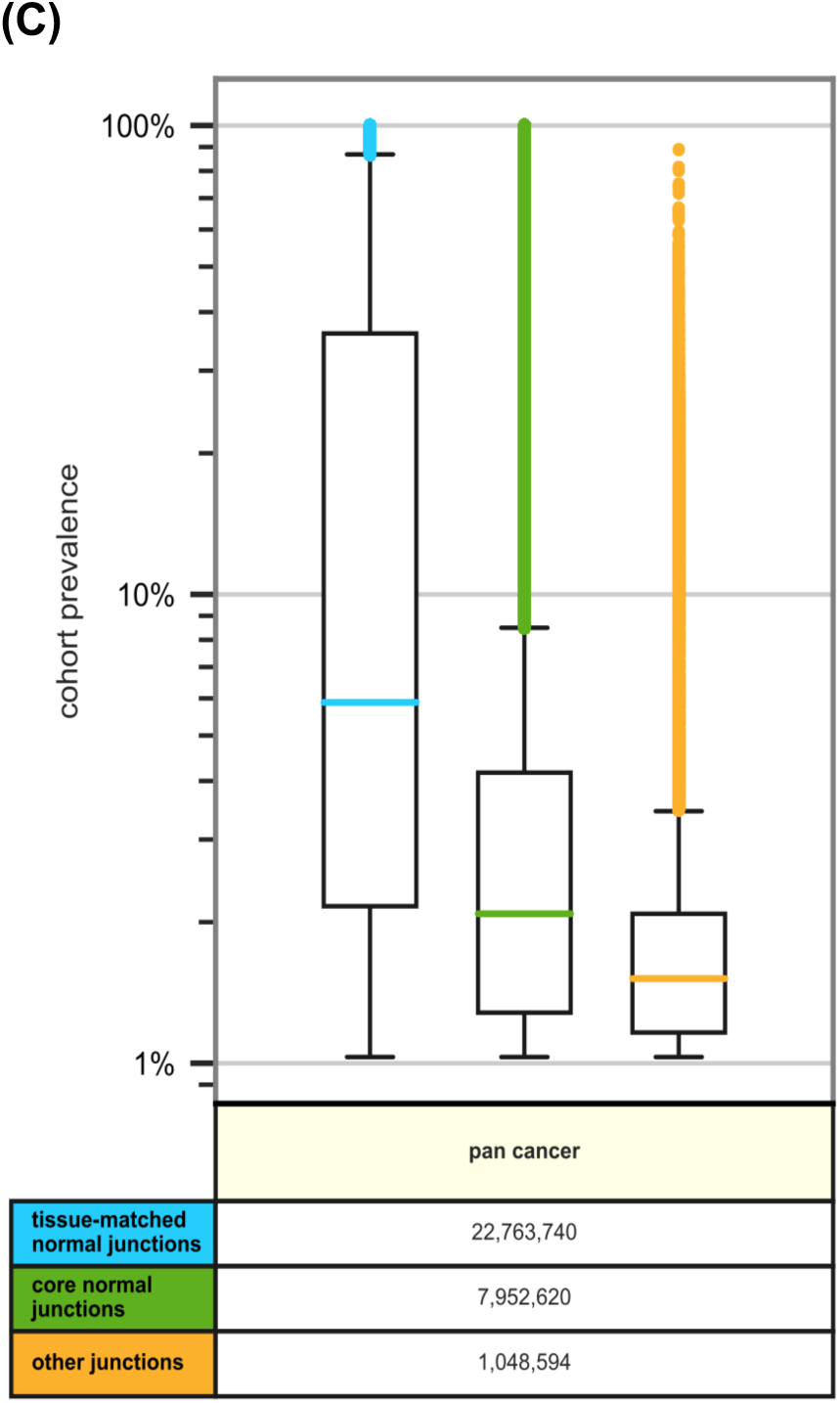
Distribution of exon-exon junctions across and within TCGA cancer cohorts. **(A)** Log-scale bar charts describing the percentage of all junctions of a given cancer-type cohort present in three sub-cohorts. Blue (left) bars give the percentage of cohort junctions found in GTEx or TCGA tissue-matched normal samples (Table S1); green (center) bars give the percentage of the remaining junctions that are found in other core normals; and yellow (right) bars give the percentage of cohort junctions found in no core normals; cancer types are ordered by relative abundance of junctions in this last set. Cancer types with no blue (left) bar have no tissue-matched normal samples (Table S1). **(B)** Log-scale sorted strip plots representing the number of non-core normals junctions per sample for each of 33 TCGA cancer types. Each point represents a single TCGA tumor sample and the width of each strip is proportional to the size of the cancer type cohort (Kahles et al., 2018). Figure S1A shows analogous data with additional filters applied. **(C)** Log-scale box plots representing the prevalences within each cancer-type cohort of junctions occurring in at least 1% of cancer-type samples, summarized across all TCGA cancer types. Junction prevalences are shown in blue (left) for those found in GTEx or TCGA tissue-matched normal samples (Table S1); junctions not present in tissue-matched normals but found in other core normals are shown in green (center); and junctions found in no core normals are shown in yellow (right). Note that any junction found in multiple cancer types is represented by multiple data points, one for each cancer type in which it is found. A detailed breakdown by TCGA cancer type is available in Figure S1D.

We next assessed the extent to which a given junction not found in core normals might be shared among multiple samples of the same cancer type. We observed that over half (52.8%) of these junctions are confined to individual samples, although a small but significant subset (0.41%) is shared across at least 5% of samples in at least one cancer-type cohort (Figure S1B). We also noted that 40.6% of novel junctions are shared between multiple cancer types, with a total of 1,609 junctions present in at least 5% of samples each across two or more TCGA cancer cohorts (Figure S1C). Sharedness was significantly higher among junctions that were also present in normal tissues (Figures 1C and 1D). We observed that the number of junctions not found in core normals per patient was comparable for patients with and without splicing factor-associated mutations across all cancer types, with the exception of breast adenocarcinoma (Figures S1E and S1F). We also observed that splicing-associated mutations had minimal effect on the sharedness within a cancer-type cohort of junctions not found in core normals (Figure S1G and S1H).

We finally assessed whether these junctions were also shared among independent cancer cohorts, using publicly available RNA-seq data in the SRA. Many TCGA cancer junctions not found in core normals were found to occur in cancer-type matched SRA samples: 11 of 14 cancer types had more than 50 junctions in common between the matched cohorts. Moreover, we found that junctions also present in matched SRA cancer cohorts were associated with significantly higher levels of sharedness in the TCGA cohort (H statistic = 3.85-2,803 and p = <0.0001-0.0495; Figure S1I).

### Shared novel junctions in cancer distinguish cancer identity and subtype

We hypothesized that a high level of exon-exon junction sharedness across samples is likely to be reflective of underlying conserved biological processes (e.g. among normal tissues). We therefore investigated the sharedness of novel junctions present in different cancer types. Interestingly, these novel junctions can readily distinguish disparate cancer types and show similarities among cancer types with shared biology, such as cutaneous and uveal melanomas (Figure 2A). These novel junctions also reflect shared biology among additional cancer types with similar anatomic origins: colon and rectal adenocarcinoma, clear cell, chromophobe, and papillary renal cell carcinomas, low and high grade gliomas, and stomach and esophageal adenocarcinomas (Figure 2A). Shared junctions from several cancer types also demonstrate similarities by histological subtype despite their differing anatomical origins, for instance squamous cell carcinomas of the lung, cervix, and head and neck (Figure 2A, Figure S2A), consistent with previously published work (Lin et al., 2017). Moreover, shared novel junctions are readily able to distinguish distinct histological subtypes of sarcoma and cervical cancer, among other diseases (Figure 2B). Using non-cancer cell types from the Sequence Read Archive (SRA) (Leinonen et al., 2011) we found that “novel” junctions from cancers arising from cell and tissue types poorly represented in GTEx normal tissue samples (e.g. melanocytes), or not present in GTEx at all (e.g. thymus tissue), can be found in many samples of the corresponding cell or tissue types of origin (Figure 2C, Table S1). Sample-to-sample comparisons of all junctions from these rare-cell type cancers also show more similarity with cell type-matched normal samples from the SRA than with bulk tissue from GTEx (Figure S2B).

**Figure 2:**
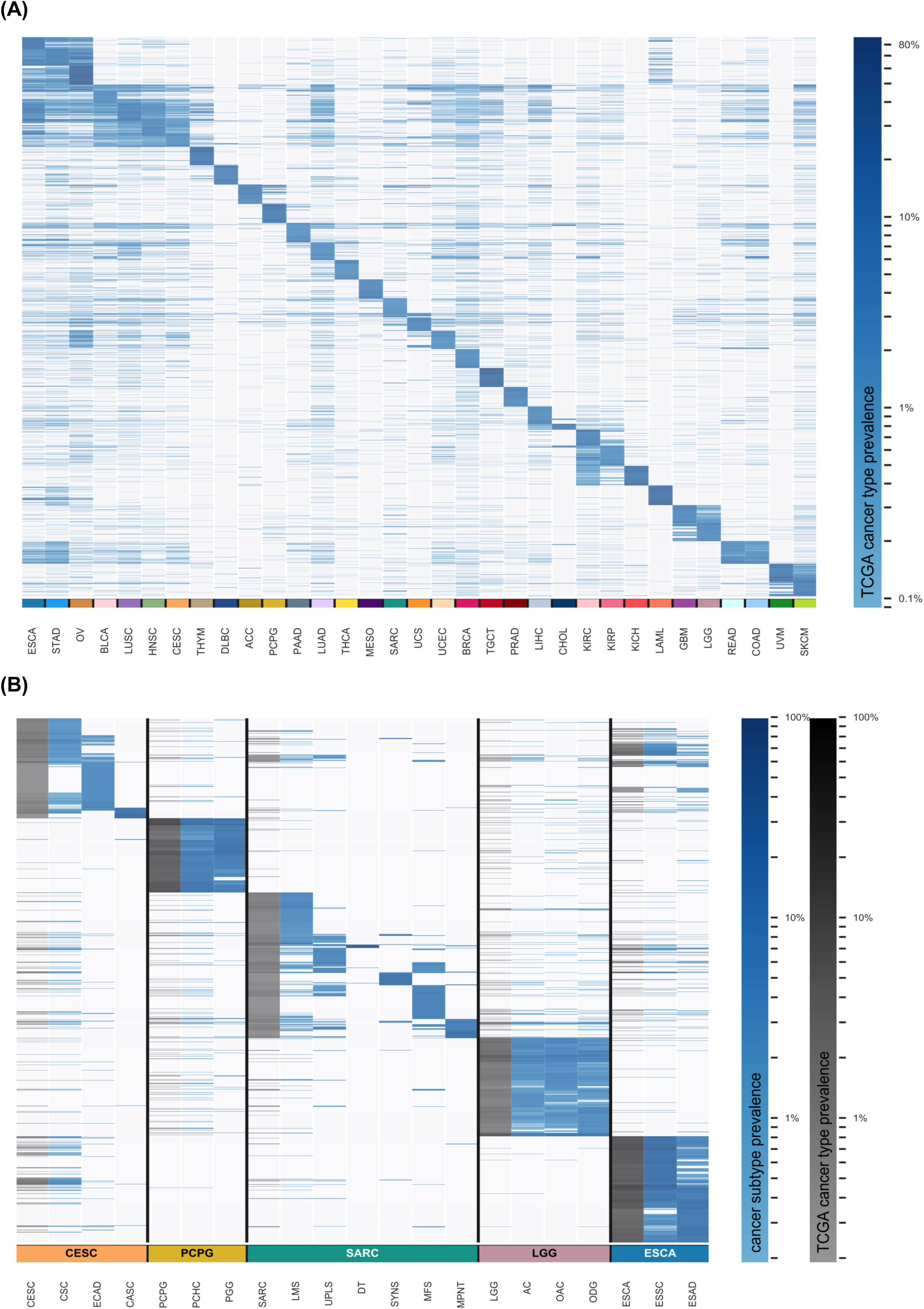
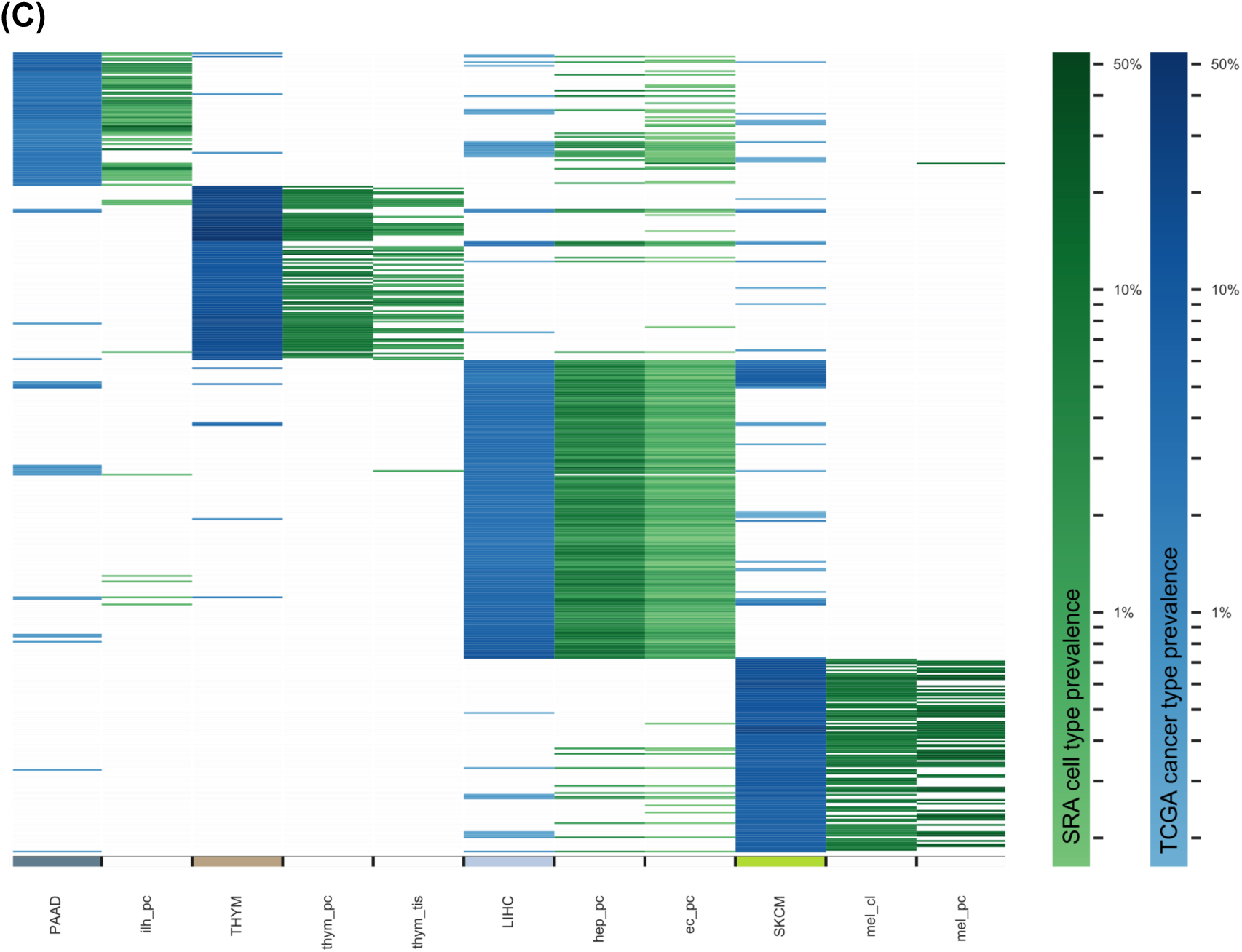
Clustering by cohort prevalence of shared novel junctions not found in core normal samples. **(A)** Heatmap showing junction prevalences across every TCGA cohort for each cancer type’s top 200 shared junctions that are at least 1% prevalent in that cancer type and are not found in any core normal samples. **(B)** Heatmap showing shared junction prevalences across selected TCGA cancer types and their assigned histological subtypes for each subtype’s top 200 shared junctions that are at least 1% prevalent in that subtype and are not found in any core normal samples. See Table S1 for TCGA subtype abbreviations. **(C)** Heatmap showing shared junction prevalences across selected TCGA cancer types and a set of their matched SRA tissue and cell types of origin, for each cancer type’s top 200 shared junctions that are at least 1% prevalent in that cancer cohort and are not found in any core normal samples. See Table S3 for SRA sample type abbreviations.

### Novel junctions in cancer are found among developmental and known cancer-related pathways

As many cancers are thought to recapitulate normal developmental pathways (Borczuk et al., 2003; Huang et al., 2009; Naxerova et al., 2008), we further hypothesized that a subset of cancer-specific junctions may reflect embryological and developmental splicing patterns. We therefore compared cancer junctions not found in core normals with those from SRA samples pertaining to zygotic, placental, embryological, and fetal developmental processes (see Methods). On average, per cancer type, 15.4% of these junctions occur in SRA developmental cell or tissue samples, and 26.5% and 2.7% occur in samples from selected SRA normal adult tissues and cell types and SRA normal stem cell samples, respectively (Figures 3A and S3A). The remaining significant majority of these cancer junctions not found in core normals were also not present in any non-cancer SRA tissue or cell type studied (64.9% on average per cancer type cohort, Figures 3A and S3A). Many of these novel “unexplained” junctions still exhibit high levels of sharedness both within (Figures S3B and S3C) and between (Figure S3D) different cancer types. At the upper end, 16 of these shared junctions were found in more than 10% of samples in each of two or more cancer types (Table S4).

**Figure 3:**
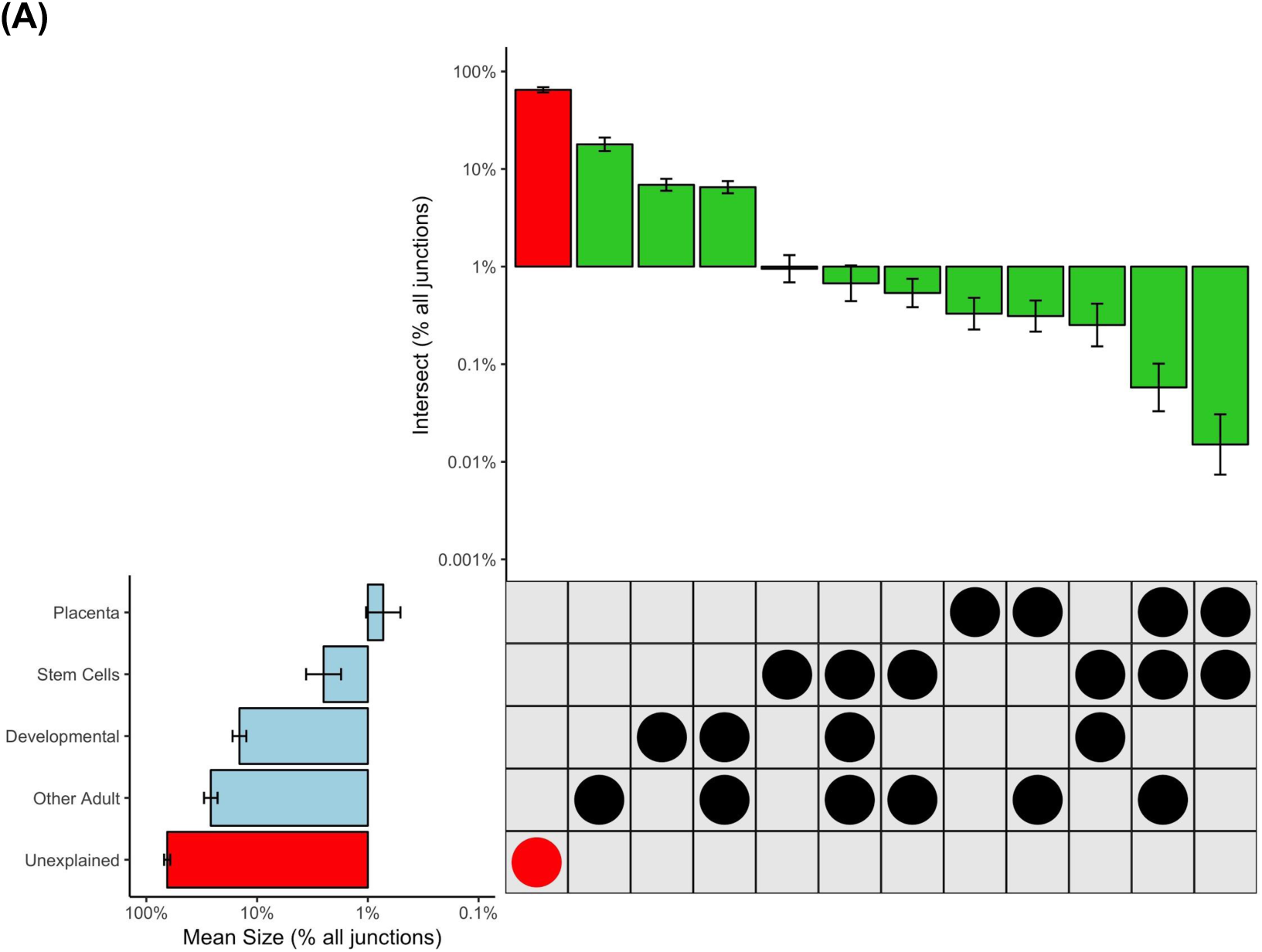
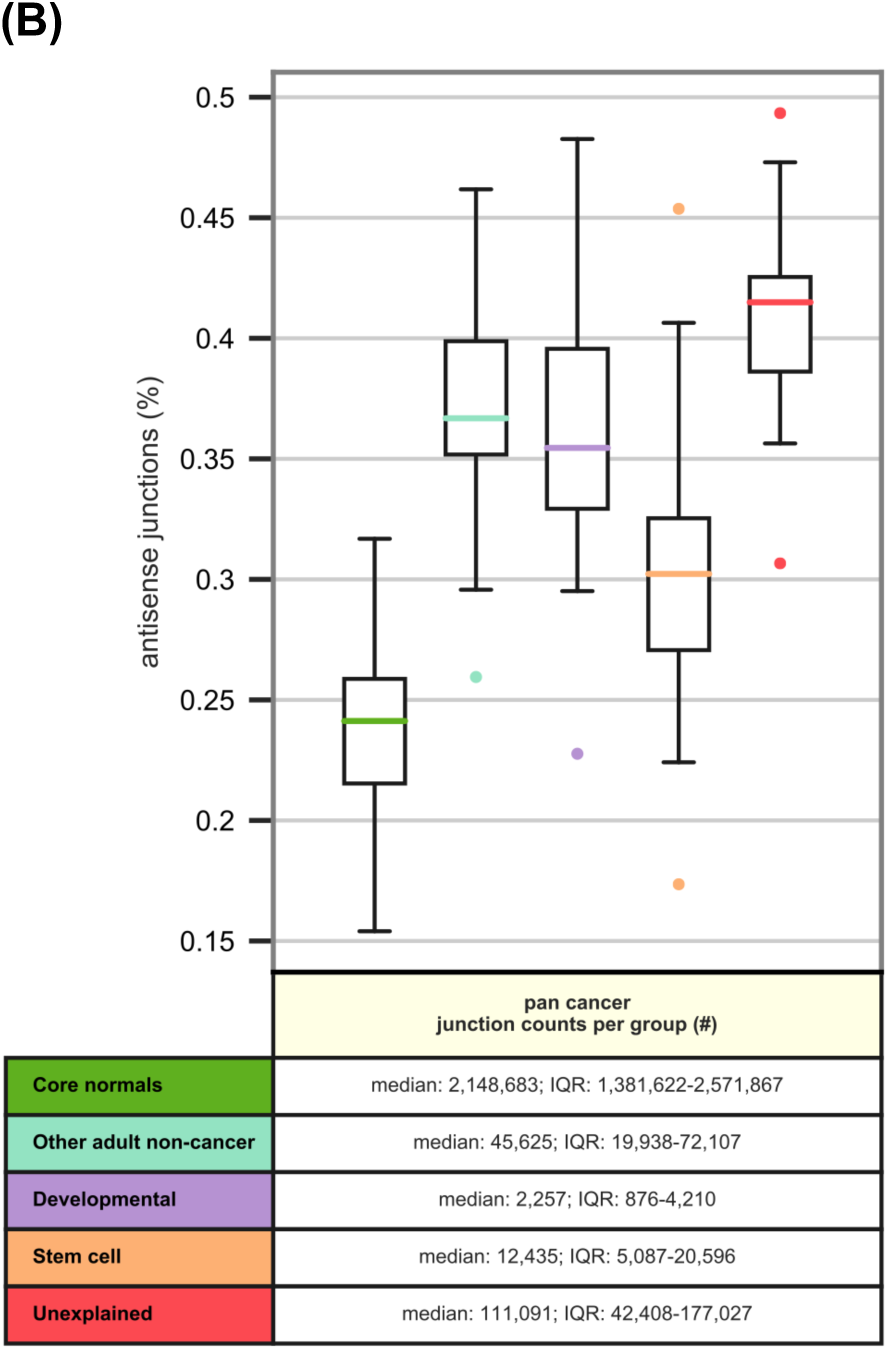
Junction set assignments and antisense junction prevalence in additional normal tissue and cell type categories from the Sequence Read Archive, across cancers. **(A)** Upset-style plot with bar plots showing junction abundances across major sets (left) and set overlaps (top) across 33 cancers (error bars). Shown junctions are absent from all core normals. Unexplained junctions (red highlights) comprise junctions not present in any set categories studied (see also expanded set assignments in Figure S3A). The developmental set comprises human development-related junctions not present in the category placenta. Scale is log10 of percent of junctions not found in core normals, calculated for each cancer. **(B)** Box plots showing the ratio of antisense-gene junctions to all junctions for (green) junctions found in core normals; (aqua) junctions not found in core normals but found in other selected non-cancer adult tissue and cell samples from the SRA; (lavender) junctions not found in core normals or SRA non-cancer adult samples but found in selected developmental samples on the SRA; (apricot) junctions not found in core normals or SRA non-cancer adult samples but found in selected stem cell samples on the SRA; and (red) junctions not found in core normals or selected non-cancer adult, developmental, or stem cell samples from the SRA.

We note that the liberal set inclusion criterion we employed may reduce our ability to identify robust cancer-specific biology among unexplained junctions. For instance, the well-described deletion causing a splicing of exons 1 and 8 (*EGFRvIII*) occurs in 29.4% of TCGA patients with glioblastoma multiforme (GBM) and in no core normals, but is also present in a single read from a single human epithelial cell line sample on SRA, and therefore is classified not as an unexplained cancer-specific junction but as “adult non-cancer.” However, this set inclusion condition does allow for the identification of some cancer-specific biology of interest. For instance, rarer alternative *EGFR* splicing events were detected in the unexplained set, such as *EGFRvIII* with an alternate exon 1 joined to exon 8 (chr7:55161631-55172981), detected in 2 patients with GBM and 1 patient with low grade glioma; the same alternate exon 1 joined with two alternate exon 16s (chr7:55161631-55168521 and chr7:55161631-55170305) (detected in 1 and 2 GBM patients, respectively); and the same alternate exon 1 joined with exon 20 (chr7:55161631-55191717) in 2 GBM patients. An alternative filtering approach that instead requires two samples per SRA category to define junction set membership yields a greater number of unexplained junctions (Table S1D and Figures S3E and S3F).

We observed a number of unexplained junctions shared by unusually large proportions of ovarian cancer (OV) samples in TCGA, including one cancer-specific junction (chr16:766903-768491 on the minus strand) present in the highest proportion of samples in any TCGA cohort (81.3%, or 350 of 430 samples in OV). This junction occurs in an antisense transcript of *MSLN*, which codes for a protein known to bind to the well-known ovarian cancer biomarker *MUC16* (CA125) (Felder et al., 2014; Kaneko et al., 2009). Another unexplained junction (chr19:8865972-8876532 on the minus strand) is in the *MUC16* region itself and is present in 42.8%, or 184 of 430 samples in OV. In all, we identified 34 cancer-specific junctions present in >40% of OV samples. We further identified several novel pan-cancer splice variants (chr16:11851406 with chr16:11820297, chr16:11821755, and chr16:11828391, each present across up to 8 different cancers) in *RSL1D1* and its neighboring *BCAR4*, a long noncoding RNA known to promote breast cancer progression (Godinho et al., 2011; Li et al., 2016).

Among all otherwise unexplained junctions, an average of 4.78% across cancer types are associated with known cancer-predisposing or cancer-relevant loci. Further, an elevated proportion of otherwise unexplained junctions (on average, 40.9%) occur in likely antisense transcripts and may therefore be of reduced interest as candidate neoantigens, but sustained interest in terms of cancer biology (Figure 3B, Table S5).

## Discussion

Previous studies have established the importance of alternative and aberrant splicing in cancer prognosis (Bjørklund et al., 2017; Li et al., 2017; Marcelino Meliso et al., 2017; Robertson et al., 2018; Zhu et al., 2018; Zong et al., 2018) and have begun to explore its potential relevance in cancer immunotherapy (Kahles et al., 2018; Wood et al., 2019). In this study, we explore “novel” exon-exon junction use among cancers with respect to a broad collection of normal tissues and cells. This is the largest such study to-date, integrating RNA-seq data from 10,549 tumor samples across 33 TCGA cancer types, 788 paired normal samples across 25 TCGA cancer types, 9,555 normal samples across 30 GTEx tissue types, and 12,231 human samples from the SRA (10,827 samples from 33 normal tissue and cell types and 1,404 samples from 14 cancer types) (Tables S1 and S2). To the best of our knowledge, this is also the first study to examine the novelty of cancer junctions from the perspective of immune tolerance, considering all adult normal tissue types as potential sources of tolerogenic peptides rather than only the closest matched normal tissues. Moreover, this is the first study to quantitatively interrogate the sharedness of novel exon-exon junctions both within and across cancer types, demonstrating that these junctions can distinguish cancers and their subtypes. We finally demonstrate that there is no one-size-fits-all definition of “novel” splicing, noting that purportedly cancer-specific junctions may in fact be present among, and perhaps biologically consistent with, a repertoire of embryological, developmentally-associated, and other cell types.

This study also has several limitations. We focus on the importance of exon-exon junctions as the predominant metric of alternative splicing. However, there are other sources of RNA variation (e.g. intron retention events (Smart et al., 2018) and RNA editing) that we do not explicitly study here, but which could be equally good sources of novel, cancer-specific protein sequence for immunotherapeutic and other applications. Additionally, there is substantial variability among analytical methods for identifying these exon-exon junctions. We note significant discordance between results of analyses of the same data using different junction filtering methods. While the same phenomena and general results appear to hold true independent of analytical technique, the identity and relative novelty of individual “cancer-specific” junctions vary between our results and those previously published (Kahles et al., 2018). We also acknowledge that GTEx and the SRA combined do not account for all sources of normal tissue(s) in the human body, and further acknowledge that the sample metadata used to search the SRA may be an imperfect surrogate for actual tissue/sample identities. Our assessment of embryological and developmentally-associated junctions is also limited by a relatively small number of relevant RNA-seq samples available on the SRA. Finally, due to the short-read nature of these RNA-seq data, we make no attempt to predict putative neoepitopes from cancer-specific junctions as we cannot confidently recapitulate reading frame or broader sequence context from isolated exon-exon junctions.

While cancer-specific exon-exon junctions may indeed be a source of neoepitopes, their sharedness across individuals and occurrence in cancer-relevant loci (e.g. *EGFR, MUC16*) are suggestive of underlying but as-of-yet unexplored biology. This sharedness does not appear to be related to variants in splicing factor or splicing-associated proteins, and is not wholly explained by recapitulation of embryological/developmental transcriptional profiles. As such, we see this work as opening a broad area of future research into the role and relevance of these novel recurring exon-exon junctions.

## Acknowledgements

We thank Paul Spellman for helpful discussions and Chris Wilks for facilitating Snaptron queries. We thank Mary Wood for her critical reading of the manuscript. The results published here are in part based upon data generated by the TCGA Research Network: https://www.cancer.gov/tcga. The Genotype-Tissue Expression (GTEx) Project was supported by the Common Fund of the Office of the Director of the National Institutes of Health, and by NCI, NHGRI, NHLBI, NIDA, NIMH, and NINDS.

## Disclaimers

The contents do not represent the views of the U.S. Department of Veterans Affairs or the United States Government.

## Author Contributions

Conceptualization, J.K.D., R.F.T., and A.N.; Methodology, J.K.D., R.F.T., and A.N.; Software, J.K.D., B.R.W., S.K.M., and A.N.; Validation, J.K.D. and A.N.; Formal Analysis, J.K.D., B.R.W., S.K.M., and A.N.; Investigation, J.K.D., B.R.W., S.K.M., and A.N.; Resources, R.F.T. and A.N.; Data Curation, J.K.D. and B.R.W.; Writing – Original Draft, J.K.D., B.R.W., R.F.T., and A.N.; Writing – Review & Editing, J.K.D., B.R.W., S.K.M., R.F.T., and A.N.; Visualization, J.K.D., B.R.W., and S.K.M.; Supervision, R.F.T. and A.N.; Project Administration, R.F.T. and A.N.; Funding Acquisition, R.F.T. and A.N.

## CONTACT FOR REAGENT AND RESOURCE SHARING

Further information and requests for resources and reagents should be directed to and will be fulfilled by the Lead Contact, Abhinav Nellore.

## METHOD DETAILS

### Data Download

Previously called exon-exon junction data including phenotype table, bed and coverage files for both TCGA and GTEx v6 were downloaded from the recount2 service at https://jhubiostatistics.shinyapps.io/recount/ (Collado-Torres et al., 2017). The metaSRA (Bernstein et al., 2017) web query form at http://metasra.biostat.wisc.edu/ was queried for experiment accession numbers for 1) non-cancer cell and tissue type samples (see Table S1 for cancer-matched samples and Table S3 for non-cancer samples, and “Selection of SRA tissue and cell types” for a description of how these samples were chosen) and 2) TCGA-matched cancer types (see Table S1). For the non-cancer samples, the term “cancer” was explicitly added as an excluded ontology term in the query, and the resulting files were filtered to remove any samples with “tumor” in the sample_name field. The resulting accession numbers were queried against the Snaptron junction database using the query snaptron tool (Wilks et al., 2018), yielding junctions for the tissue and cell types of interest that were downloaded. Patient somatic mutation calls were downloaded from the GDAC firehose (Broad Institute TCGA Genome Data Analysis Center, 2016), while a list of human splicing-associated gene mutations (keyword search “mRNA splicing [KW-0508]”) was downloaded from the UniProt database (UniProt Consortium, 2019). Two lists of cancer-associated genes were downloaded: the COSMIC cancer gene census cancer gene list from https://cancer.sanger.ac.uk/census (Sondka et al., 2018), and the OncoKB cancer gene list from https://oncokb.org/cancerGenes (Chakravarty et al., 2017).

### Indexing of GTEx and TCGA junctions

The GENCODE gene transfer format (.gtf) file was parsed to collect full coordinates and left and right splice sites of junctions from annotated transcripts and a searchable tree of protein-coding gene boundaries. The GTEx phenotype file was parsed to collect tissue of origin information and donor gender; bone marrow samples derived from leukemia cell line cells were eliminated. The TCGA phenotype file was parsed to collect information on cancer type, cancer stage at diagnosis, patient gender, vital status, and sample type (primary tumor, matched normal sample, recurrent tumor, or metastatic tumor). Cancer subtype classifications were collected for five cancer types beyond their TCGA designations (Figure 2B, Table S1): cervical squamous cell carcinoma and endocervical adenocarcinoma was separated into cervical squamous cell carcinoma, endocervical adenocarcinoma, and cervical adenosquamous; esophageal carcinoma was separated into esophagus adenocarcinoma and esophagus squamous cell carcinoma; brain lower grade glioma was separated into astrocytoma, oligoastrocytoma, and oligodendroglioma; sarcoma was separated into leiomyosarcoma, myxofibrosarcoma, malignant peripheral nerve sheath tumors, desmoid tumors, dedifferentiated liposarcoma, synovial sarcoma, and undifferentiated pleomorphic sarcoma; and pheochromocytoma and paraganglioma were separated. A new SQLite3 database was created to index all GTEx and TCGA junctions, with linked tables containing 1) sample ids and associated junction ids; 2) sample ids and phenotype information for each sample; and 3) junction ids and junction information including 0-based closed junction coordinates, GENCODE annotation status, and location within protein coding gene boundaries. SQL indexes were created on junction ID and sample ID columns for fast and flexible querying.

### Selection of cancer-specific junctions

For all analyses we apply a light filter, requiring a junction to have at least a two-read coverage across GTEx, TCGA, and the selected cancer and non-cancer SRA samples, to exclude false positive junctions but allow for the existence of splicing noise. To characterize junction novelty in cancer with respect to normal cells, we defined a hierarchical filter that specifies inclusion and exclusion of junctions in different RNA-seq datasets (Table 1). In order from most to least permissive, these filters are: 1) junctions not found in tissue-matched GTEx or TCGA normal samples, 2) junctions not found in any GTEx or TCGA normal (“core normal”) samples, and 3) junctions not found in any core normal samples or in selected SRA tissue and cell type non-cancer samples. For our analyses, we do not explicitly filter on whether a junction is annotated in GENCODE. We do not set a limit on presence in the core normal sample cohorts: any junction present at any coverage level in only one sample is counted as “in” these cohorts. This yields a more stringent filter on normality than that used by the TCGA splicing paper, which uses the term “neojunctions” to refer to junctions not found in tissue-matched GTEx or TCGA normal samples, with a 10-read coverage requirement in TCGA, and allowing through the filter lowly expressed junctions in GTEx tissue-matched samples (Kahles et al., 2018).

**Table 1:** Junction novelty specification.

| Junction novelty stage | Definition |
| --- | --- |
| 0 | All junctions |
| 1+ | Junctions not found in tissue-matched GTEX or TCGA normal samples |
| 2+ | Junctions not found in any GTEX or TCGA normal (“core normal”) samples |
| 3+ | Junctions not found in any core normal samples or in selected SRA non-cancer samples |

We queried the junction database to extract junctions of interest, specifically 1) all junctions for all tumor samples of each cancer type and 2) all junctions not present in any core normal samples for each cancer type cohort, with their cohort prevalence levels. All junctions are presented in a 0-based closed coordinate system. Protein coding region presence was determined for all junctions, with location assessment as follows: the junction is categorized as protein-coding if it is present in a protein-coding gene region (with at least one junction splice site within the gene boundaries) and antisense if it is present on the reverse strand of a protein-coding gene region, based on gene regions described in GENCODE v.28 (Frankish et al., 2019). Cancer-associated genes were collected from the OncoKB and the COSMIC cancer gene census; any gene listed in one or both lists was categorized as a cancer-associated gene. Any junction assigned to a protein-coding gene region corresponding to one of these genes was categorized as associated with cancer-relevant loci.

For comparison between cancer-sample junctions found vs. not found in core normal samples, we performed a Kruskal-Wallis H-test to determine the significance of the decreased sharedness levels, since the junction prevalence data is not normally distributed and there are many fewer cancer-specific junctions than junctions found in core normal samples.

### Comparison with SRA tissue and cell types

Non-cancer sample types from the SRA were chosen via manual curation informed by a clustering of junctions according to ontology term prevalence, with commonly occurring terms that do not meaningfully distinguish junctions eliminated. The selected sample types in Table S3 comprise all non-cancer data from the SRA analyzed. All junctions for samples associated with these cell and tissue types but not with “cancer” were downloaded via Snaptron, translated to a 0-based closed coordinate system, and compared with those found in TCGA cancer samples. Junctions present in a TCGA cancer-type cohort and SRA samples from a specific assigned category determined set assignments, which were used for subsequent data analysis. To exclude false positive junctions but allow for the existence of splicing noise, only junctions with at least two reads across GTEx, TCGA, and the selected cancer and non-cancer SRA samples are considered true junctions. All SRA junctions not found in TCGA cancer samples were ignored. For the supplementary 2-sample minimum filter analysis, we retained all junctions that are present in only 1 SRA sample, but required at least two samples across the broad SRA category (adult, developmental, or stem cell) for inclusion in that set. (For developmental subsets, only one sample within a subset category was required, as long as the 2-sample criterion across the full developmental category was met.)

For comparison between TCGA cancer-sample junctions not found in core normal samples with SRA junctions from matched cancer type samples, we performed a Kruskal-Wallis H-test to determine the significance of the increased sharedness levels, since the junction prevalence data is not normally distributed and the difference in junction counts between the two cohorts (TCGA junctions in or not-in the SRA matched cohort) is large.

### Splicing Factor Mutation Analysis

Patient somatic mutation call files were downloaded from the GDAC firehose (http://gdac.broadinstitute.org/) and all silent mutations were removed. Using the remaining mutation calls, patients were classified based on two-different separation criteria: 1) whether or not they had at least one mutation in a gene that codes for a protein annotated as involved in mRNA splicing, based on the UniProt protein annotation database, and 2) whether or not they had at least 1 mutation in a gene previously identified as sQTL associated (U2AF1, SF3B1, TADA1, PPP2R1A, and/or IDH1) in the TCGA cohort by the TCGA splicing paper (Kahles et al., 2018). For each cancer type, and each stratification method, the number of cancer-specific junctions per patient was compared for patients with and without at least one mutation in the defined set (Figures S1E and S1F). Differences in the number of novel junctions across cancer types and stratification groups was assessed via two-way ANOVA with a Benjamini-Hochberg p-value correction.

In addition to comparing the levels of cancer specific junctions between patients with and without splicing associated mutations, we also compared junction sharedness based on the same two stratification criteria used above. For each cancer type, all junctions identified in two or more patients were selected. For each, the number of junction occurrences in patients with mutations in splicing associated genes was calculated and compared to the overall number of occurrences in the corresponding cancer cohort, using a Fisher’s exact test (Figures S1G and S1H).

## DATA AND SOFTWARE AVAILABILITY

All data is publicly available and accessible online. Python code and corresponding descriptors for the implementation of methods as described is publicly available on GitHub (https://github.com/JulianneDavid/shared-cancer-splicing).

## Supplemental Information

**Figure S1:**
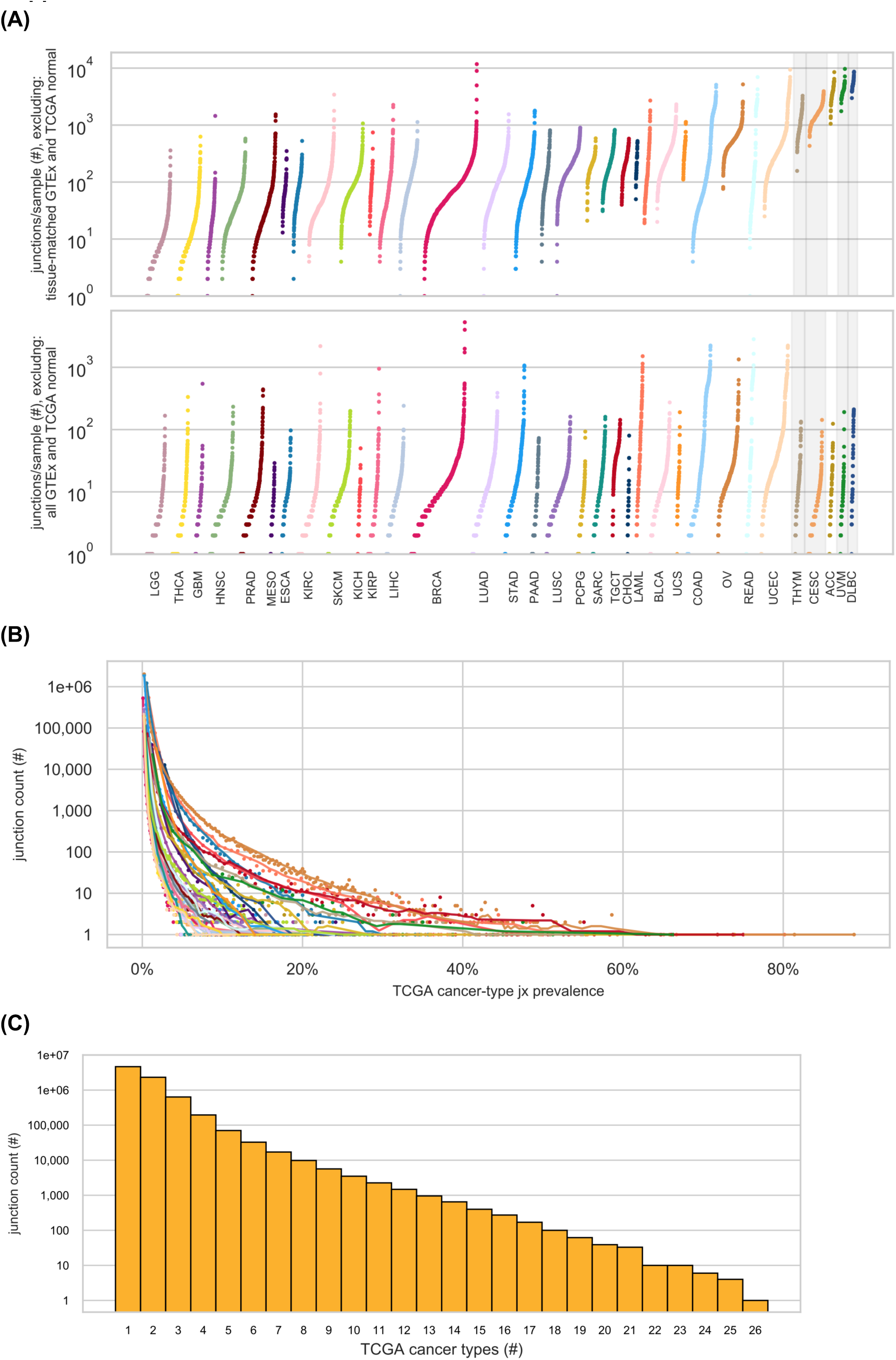
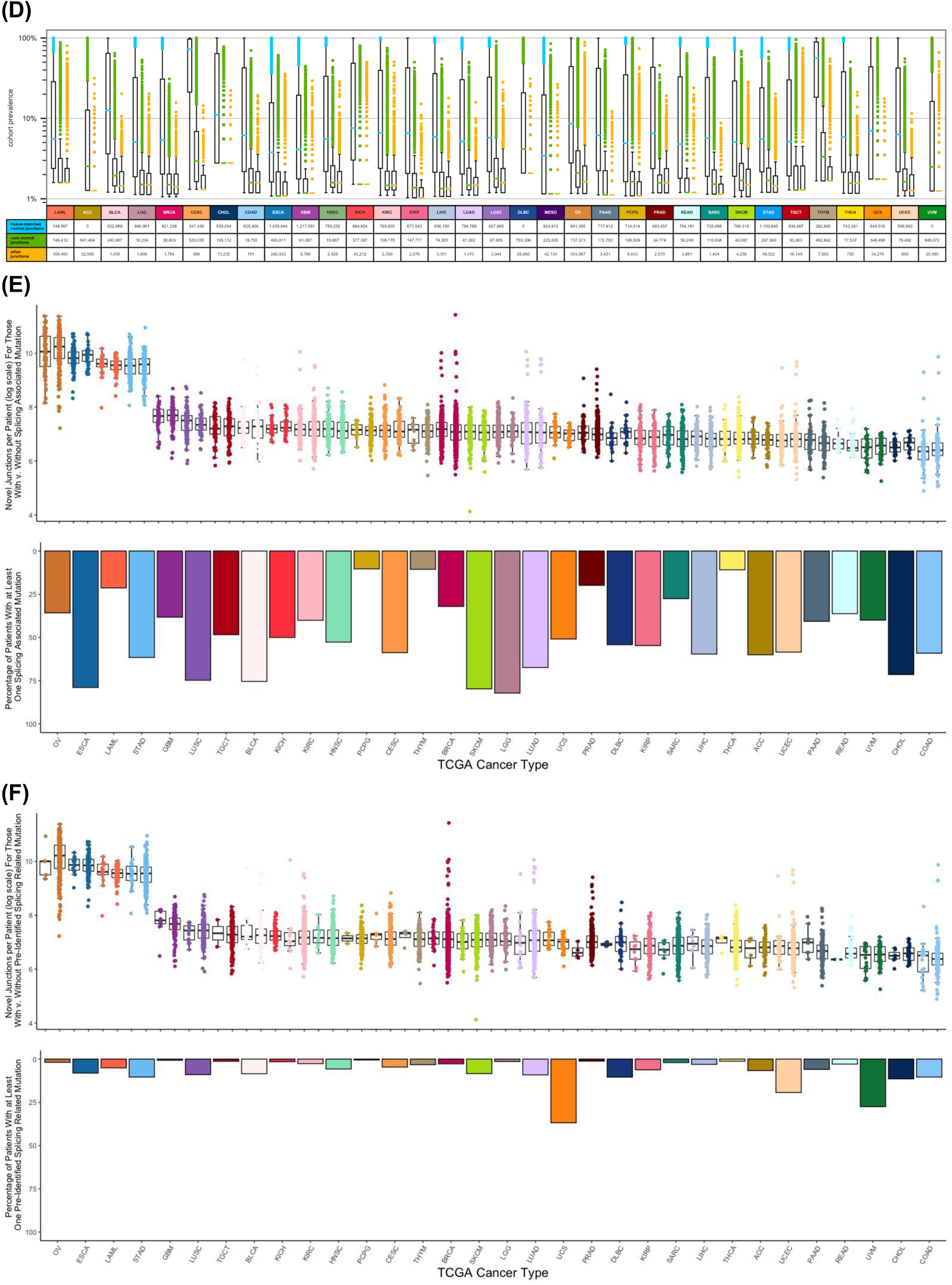
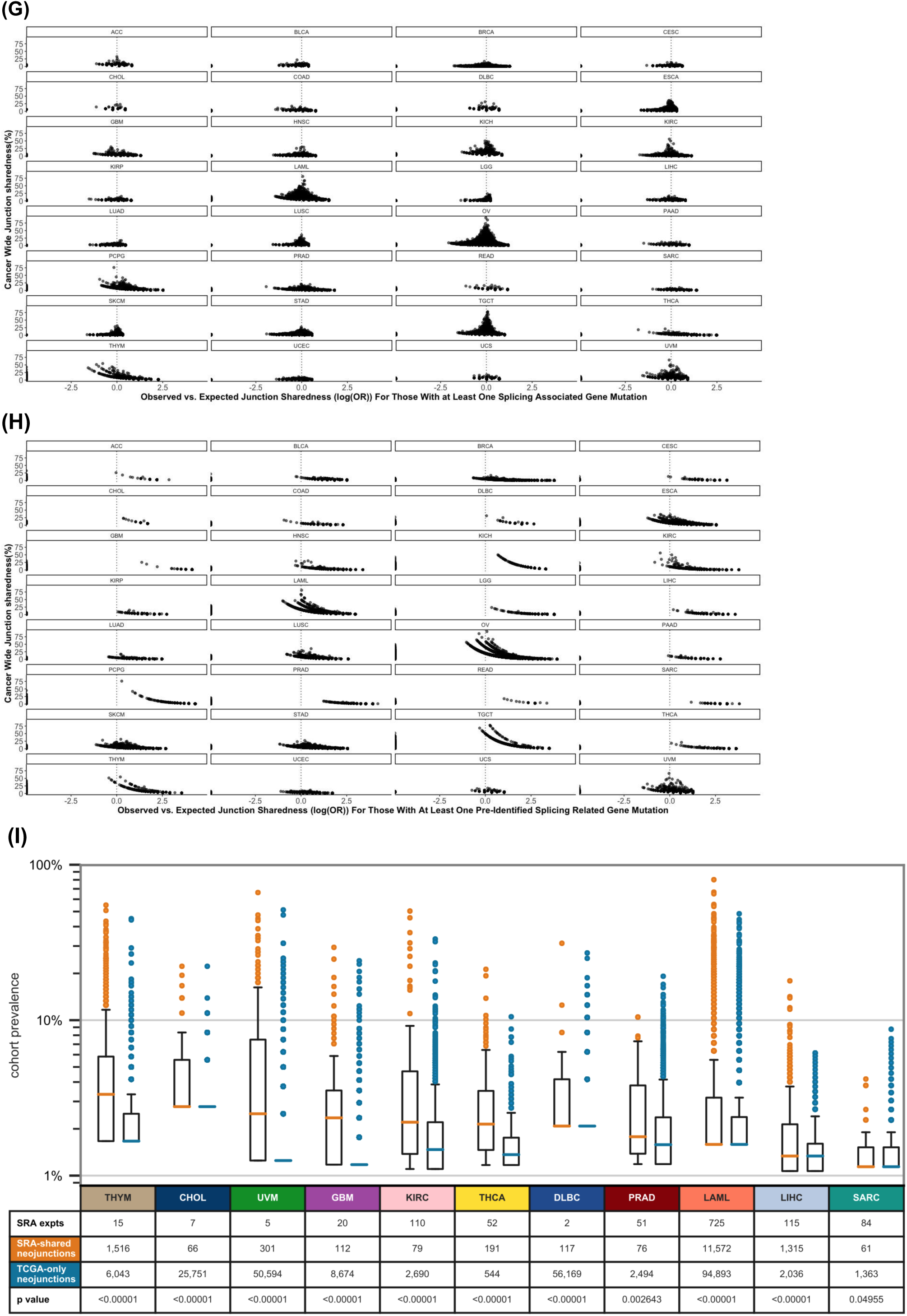
Distribution and prevalences TCGA cancer sample junctions. **(A)** Log-scale sorted strip plots representing the number of high-support junctions per sample for each of 33 TCGA cancer types where each point is a single TCGA tumor sample and the width of each strip is proportional to the size of the cancer type cohort (Kahles et al., 2018). High support requires junction coverage within a sample to be equivalent to at least 5 out of 100 million 100-base pair reads. The upper panel counts junctions not found in GENCODE annotation or in tissue-matched normal GTEx or TCGA samples (Table S1); the lower panel counts junctions not found in GENCODE annotation or in any core normal samples, differing from Figure 1B in that here, GENCODE-annotated junctions are also removed and the coverage filter is applied. The gray bars highlight TCGA cancer types with no or few tissue-matched normal samples (Table S1); note that there are orders of magnitude fewer GENCODE-annotated junctions than junctions found in tissue-matched normal samples, partially explaining the high values for THYM, CESC, UVM, and DLBC in the upper panel. **(B)** Log-scale scatter plot showing, for 33 TCGA cancer types, the number of junctions shared within each cancer-type cohort at each prevalence level, counting only junctions not found in any core normal samples. TCGA cancer type colors are as specified in Figures 1B, 2A, and S1A. Of interest with significant intra-cohort junction sharedness are, among others, ovarian carcinoma (tan), leukemia (pink), testicular germ cell tumors (red), and uveal melanoma (dark green). **(C)** Log-scale histogram showing inter-cancer sharedness of junctions not found in core normal samples. Most junctions occur in only one cancer type, but many are shared between 2 or more. **(D)** Log-scale box plots representing, for all TCGA cancer types individually, the prevalences within each cancer-type cohort of junctions occurring in at least 1% of cancer-type samples, separated into prevalences for (blue, left) junctions found in GTEx or TCGA tissue-matched normal samples (Table S1); (green, center) junctions not found in tissue-matched normals but found in other core normal samples; and (yellow, right) junctions found in no core normal samples. Any junction found in multiple cancer types is represented by multiple data points, one for each cancer type in which it is found. Figure 1C condenses all data from this figure into one pan-cancer set. **(E)** Cancer specific splicing junctions in patients with and without splicing associated gene mutations: log count of junctions not found in any core normal samples for each patient are plotted (top) across each cancer type in TCGA. Within each cancer, boxplots represent either patients with mutations in UniProt annotated splicing-related genes (left) or patients without any related mutations (right). Overall prevalence of relevant mutations in splicing-related for each cancer type are plotted below. **(F)** Presents data in the same manner as E, but with comparison between patients with mutations only in genes described in the TCGA splicing paper (Kahles et al., 2018) vs all other patients. **(G)** Analysis of junction sharedness for patients within mutational cohorts: for each cancer, junctions not found in core normal samples are plotted based on sharedness across all patients in the cancer cohort (Y-axis) compared to deviation from expected shareness (estimated odds ratio (log scale) based on Fisher’s exact test) for only patients with mutations in UniProt annotated splicing-related genes (x-axis). **(H)** Presents data in the same manner as G, but with deviation from expected sharedness calculated only for patients with mutations in genes described in the TCGA splicing paper (Kahles et al., 2018). We found that no specific junctions show significantly enriched sharedness in patients carrying relevant mutations (Fisher’s exact test FDR > 0.05 for all in both A and B), however there is a consistent shift towards higher than expected sharedness across the majority of cancers for patients carrying at least one of the mutations defined by the TCGA splicing paper (Kahles et al., 2018). **(I)** Comparison of TCGA-cohort prevalence of junctions occurring vs. not occurring in SRA cancer samples: log-scale box plots representing, for selected TCGA cancer types, the prevalences within each cancer-type cohort of junctions occurring in at least 1% of cancer-type samples, separated into prevalences for junctions (orange, left) found or (blue, right) not found in type-matched cancer sample(s) from the SRA. Selected TCGA cancer types are those for which cancer-matched SRA sample junctions are available from Snaptron (Wilks et al., 2018) and at least 50 TCGA cancer junctions not found in core normal samples are present in the cancer-type matched SRA samples. Most junctions are TCGA-specific, but junctions that are also found in an type-matched SRA cancer cohort have on average higher TCGA-cohort prevalences.

**Figure S2:**
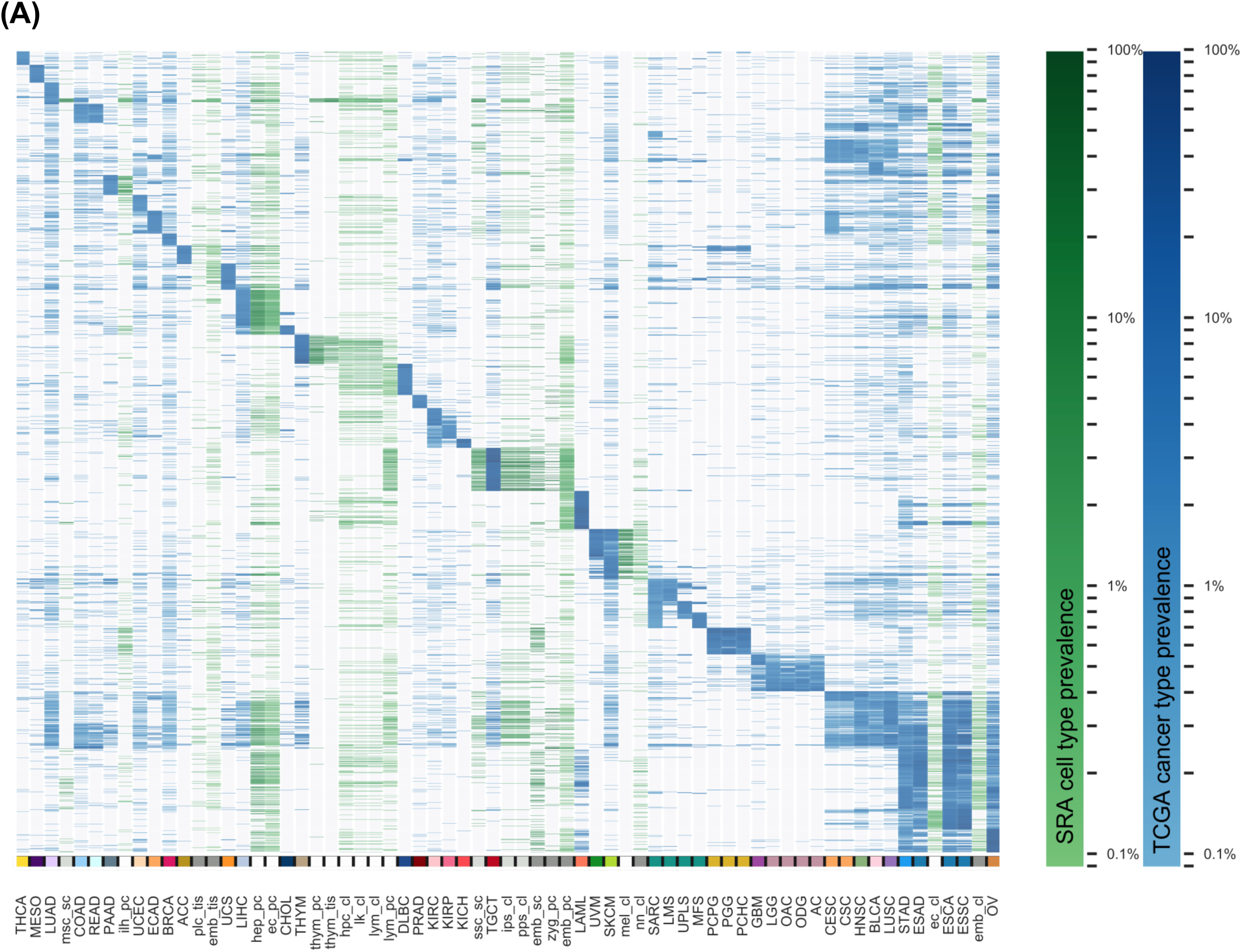
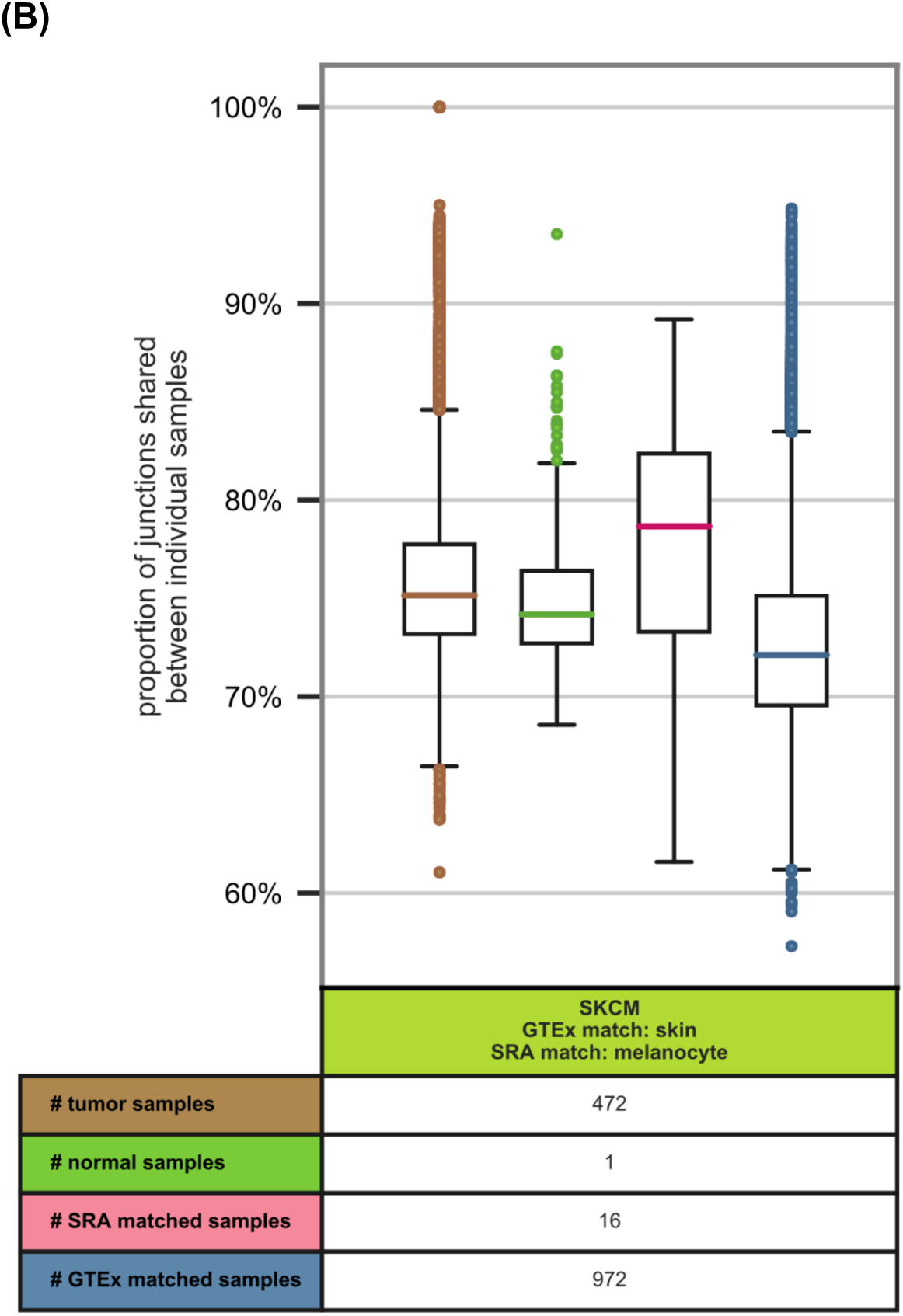
Similarity of TCGA junctions and non-cancer SRA junctions. **(A)** Clustering by cohort prevalence of junctions not found in core normal samples: heatmap showing shared junction prevalences across all TCGA cancer type and associated histological subtypes with at least 20 samples. The clustered junctions are the 200 most prevalent junctions of each cancer type or subtype that are at least 1% prevalent in that subtype and are not found in any core normal samples but are found in at least one of the 22 non-cancer tissue and cell type SRA samples. Each heatmap row represents a junction’s prevalence in each of the TCGA and SRA sample-type cohorts. The colorbar beneath the plot shows SRA tissue and cell types colored according to their assigned categories (Table S3), where white represents adult normal samples, light gray represents stem cell saamples, and darker gray represents developmental samples. **(B)** Samplewise comparison of junctions from TCGA melanoma samples and select normal samples: boxplots showing the percent of junctions shared for every pairwise combination of TCGA melanoma tumor samples with (brown) TCGA melanoma tumor samples, (grass green) the single TCGA melanoma paired normal sample, (pink) SRA normal melanocyte samples (see Tables S1 and S3), and (blue) GTEx normal skin samples. The percent of junctions shared between two samples is given by % *shared = (set A & set B) / min(len(set A), len(set B))*, where a set comprises all junctions identified in the single cancer or normal sample. TCGA melanoma cancer samples have on average a greater similarity of junctions to SRA normal melanocyte samples than to GTEx or TCGA bulk skin normal samples.

**Figure S3:**
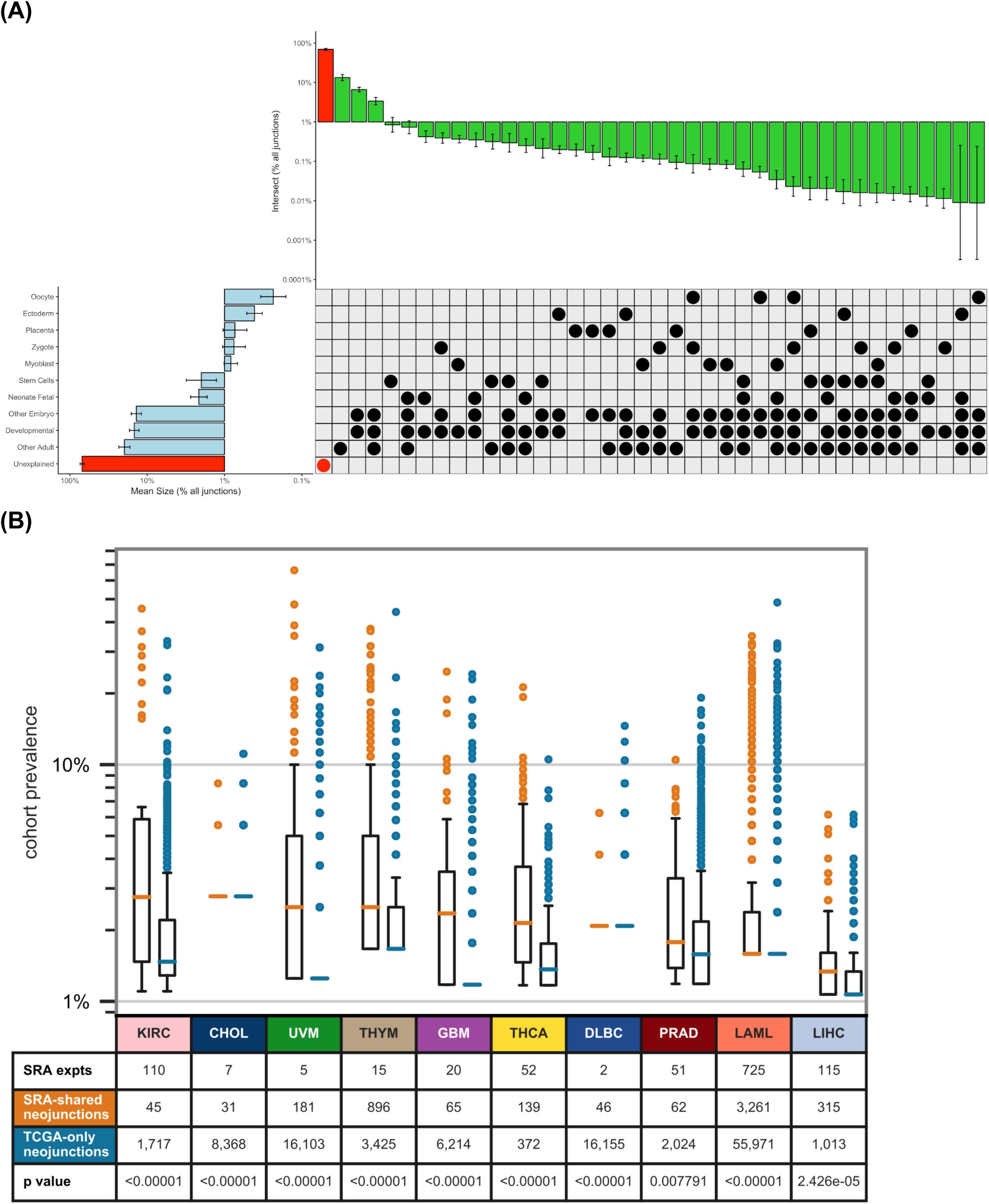
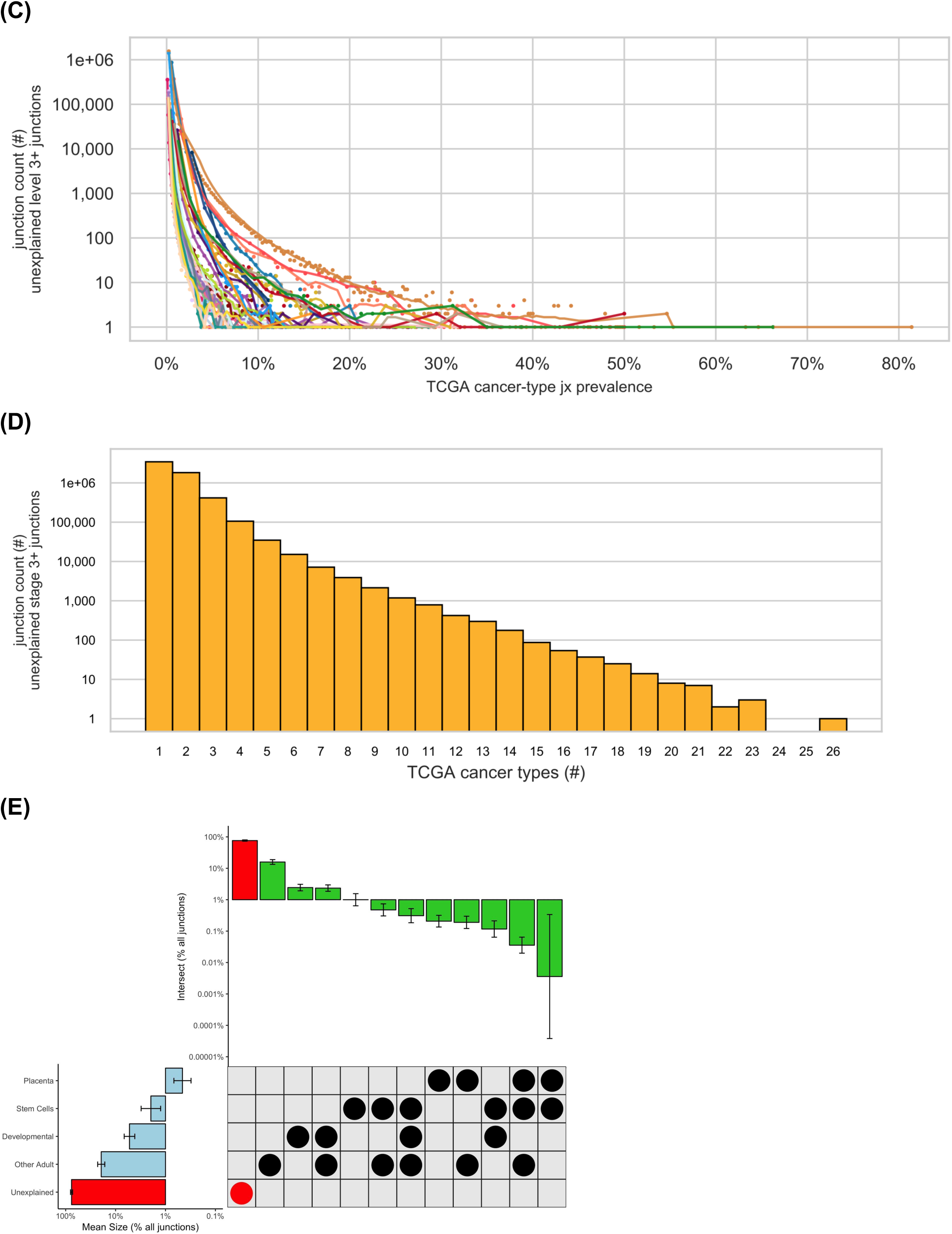
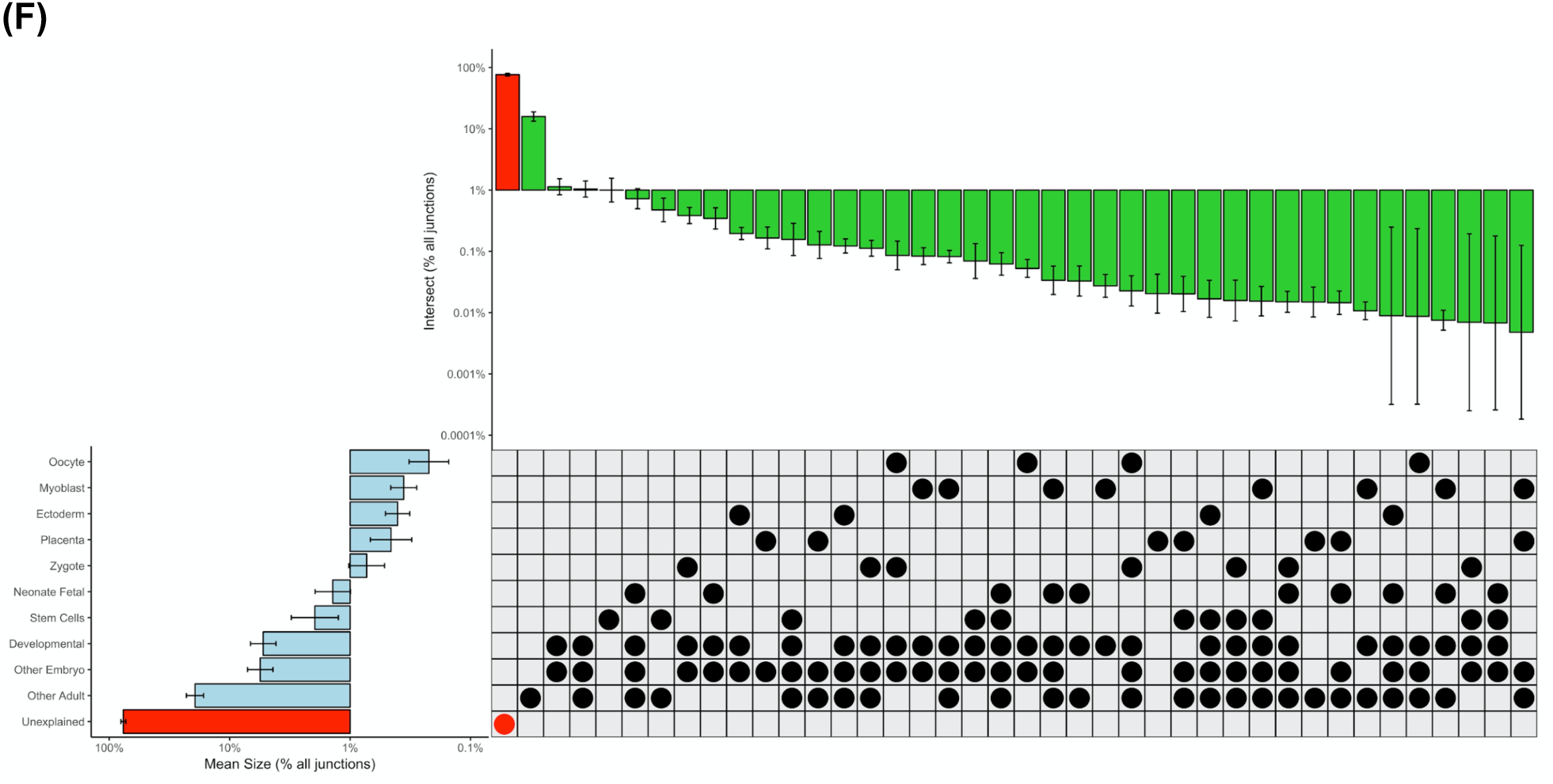
Distribution of junctions not found in core normal samples and unexplained junctions. **(A)** Expanded junction set assignments in normal tissue and cell type categories from the Sequence Read Archive, across cancers: upset-style plot with bar plots showing junction abundances across major sets and subsets (left) and set overlaps (top) across 33 cancers (error bars). Shown junctions are absent from all core normals. Unexplained junctions (red highlights) comprise junctions not present in any set categories studied (see also Figure 3A). The developmental set comprises human development-related junctions not present in the category placenta. Scale is log10 of percent of junctions not found in core normals, calculated for each cancer. **(B)** Analysis of inter- and intra-cancer sharedness of stage-3+ “unexplained” junctions: log-scale box plots as in Figure S1I but including only stage-3+ unexplained junctions not found in core normal samples or selected SRA normal adult, developmental, or stem cell samples (Table 1). Plots are presented for TCGA cancer types for which cancer-matched SRA sample junctions are available from Snaptron (Wilks et al., 2018) and at least 30 unexplained junctions are present in the cancer-type matched SRA samples. Prevalences are given within each cancer-type cohort of junctions occurring in at least 0.5% of cancer-type samples, separated into prevalences for junctions (orange, left) found or (blue, right) not found in type-matched cancer sample(s) from the SRA. For all cancer types except DLBC, most junctions are TCGA-specific, but junctions that are also found in a type-matched SRA cancer cohort have on average higher TCGA-cohort prevalences. **(C)** Log-scale scatter plot showing, for 33 TCGA cancer types, the number of level-3+ unexplained junctions shared within each cancer-type cohort at each prevalence level as in figure S1B. TCGA cancer type colors are as specified in Figures 1B, 2A, and S1A. Again, among others, ovarian carcinoma (tan), leukemia (pink), testicular germ cell tumors (red), and uveal melanoma (dark green) have significant intra-cohort junction sharedness. **(D)** Log-scale histogram showing inter-cancer sharedness of stage-3+ unexplained junctions as in Figure S1C. Again, most junctions occur in only one cancer type, but many are shared between 2 or more. **(E)** Upset-style plot with bar plots showing junction abundances across major sets (left) and set overlaps (top) across 33 cancers (error bars); similar to Figure 3A, but presence in 2 samples across the SRA sample-type category is required for inclusion in a set. Shown junctions are absent from all core normals. Unexplained junctions (red highlights) comprise junctions not present in any set categories studied. The developmental set comprises human development-related junctions not present in the category placenta. Scale is log10 of percent of junctions not found in core normals, calculated for each cancer. **(F)** Upset-style plot with bar plots showing junction abundances across major sets and subsets (left) and set overlaps (top) across 33 cancers (error bars); similar to Figure 3A, but presence in 2 samples across the SRA sample-type category is required for inclusion in a set. Shown junctions are absent from all core normals. Unexplained junctions (red highlights) comprise junctions not present in any set categories studied. The developmental set comprises human development-related junctions not present in the category placenta. Scale is log10 of percent of junctions not found in core normals, calculated for each cancer.

**Table S1:**
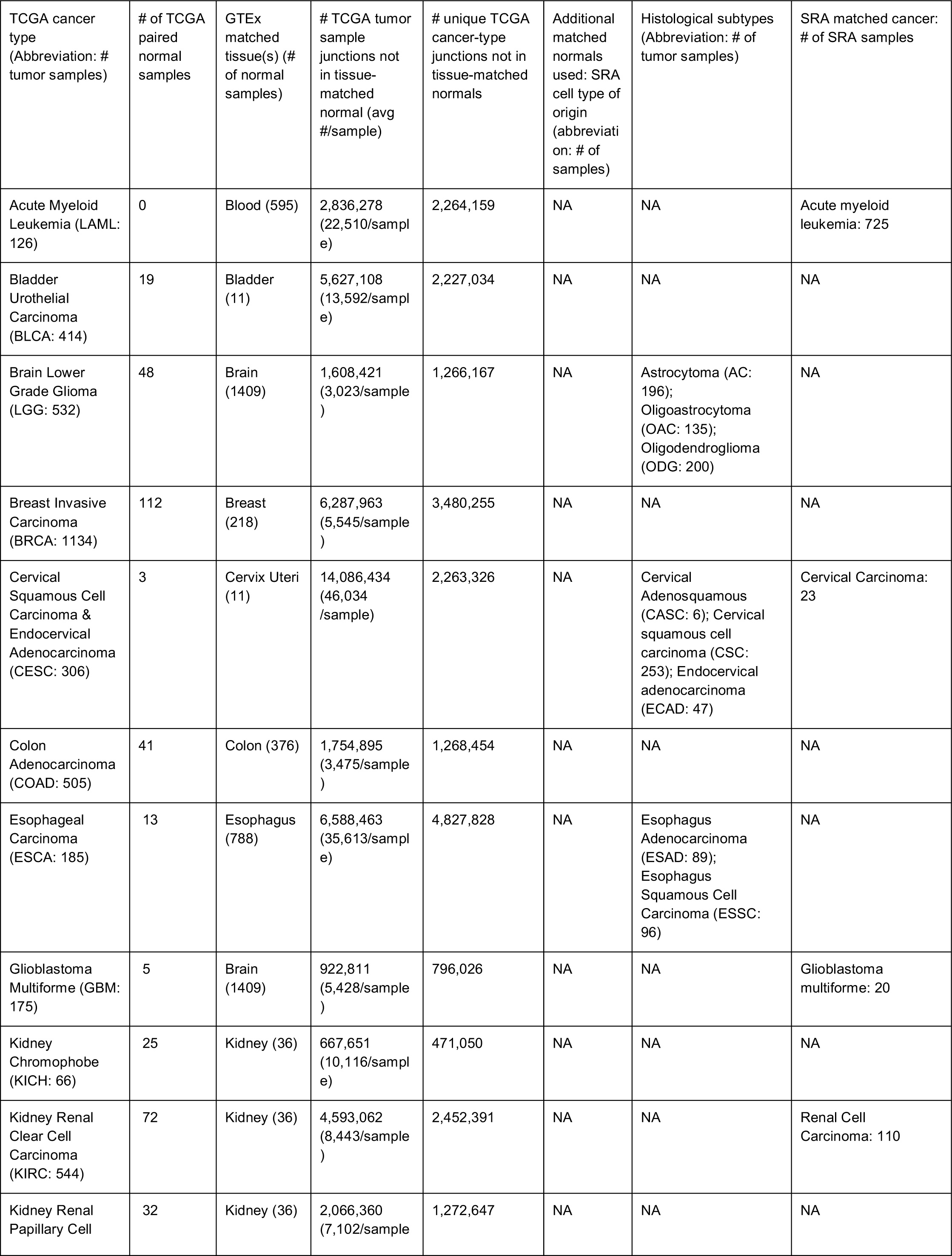
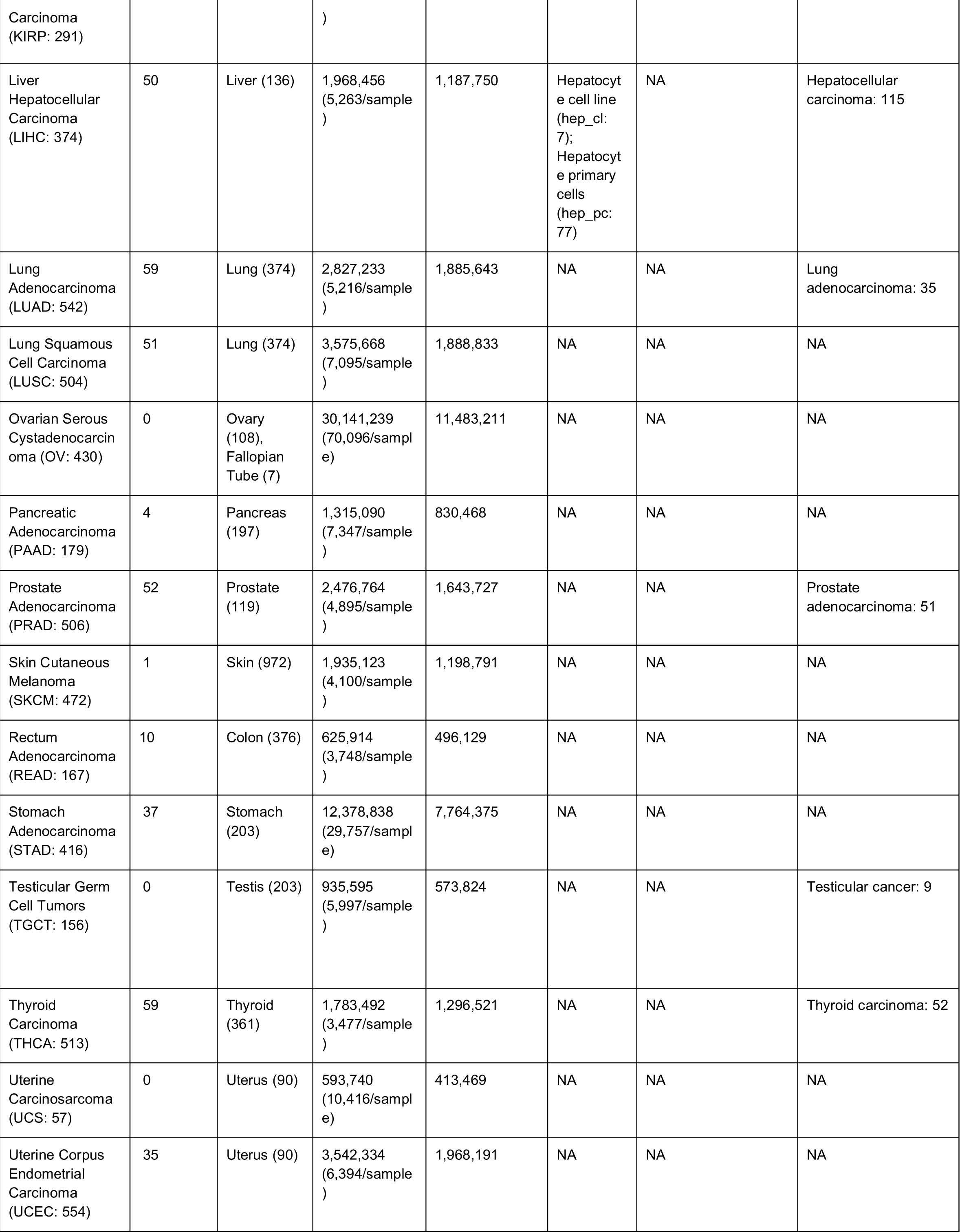
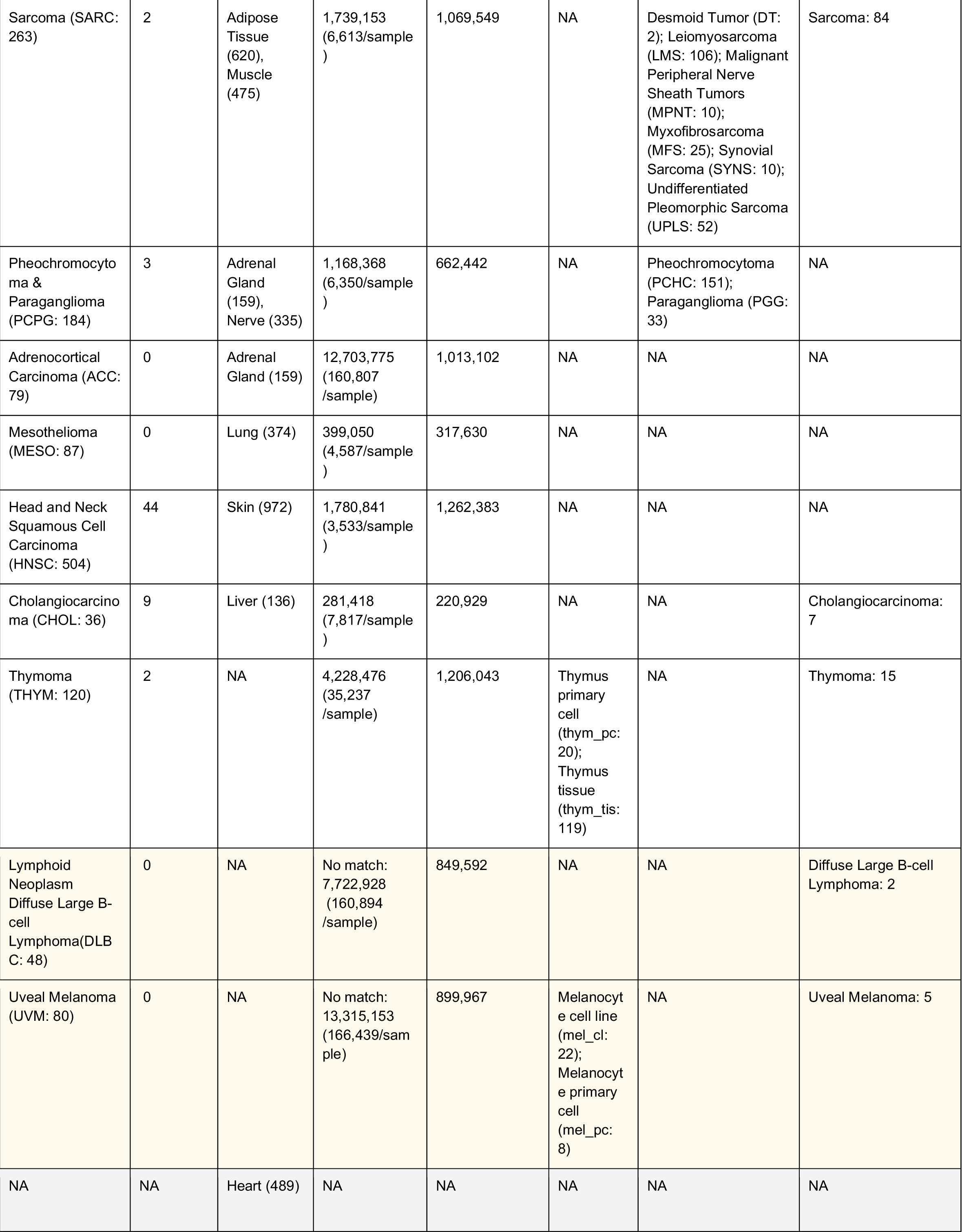
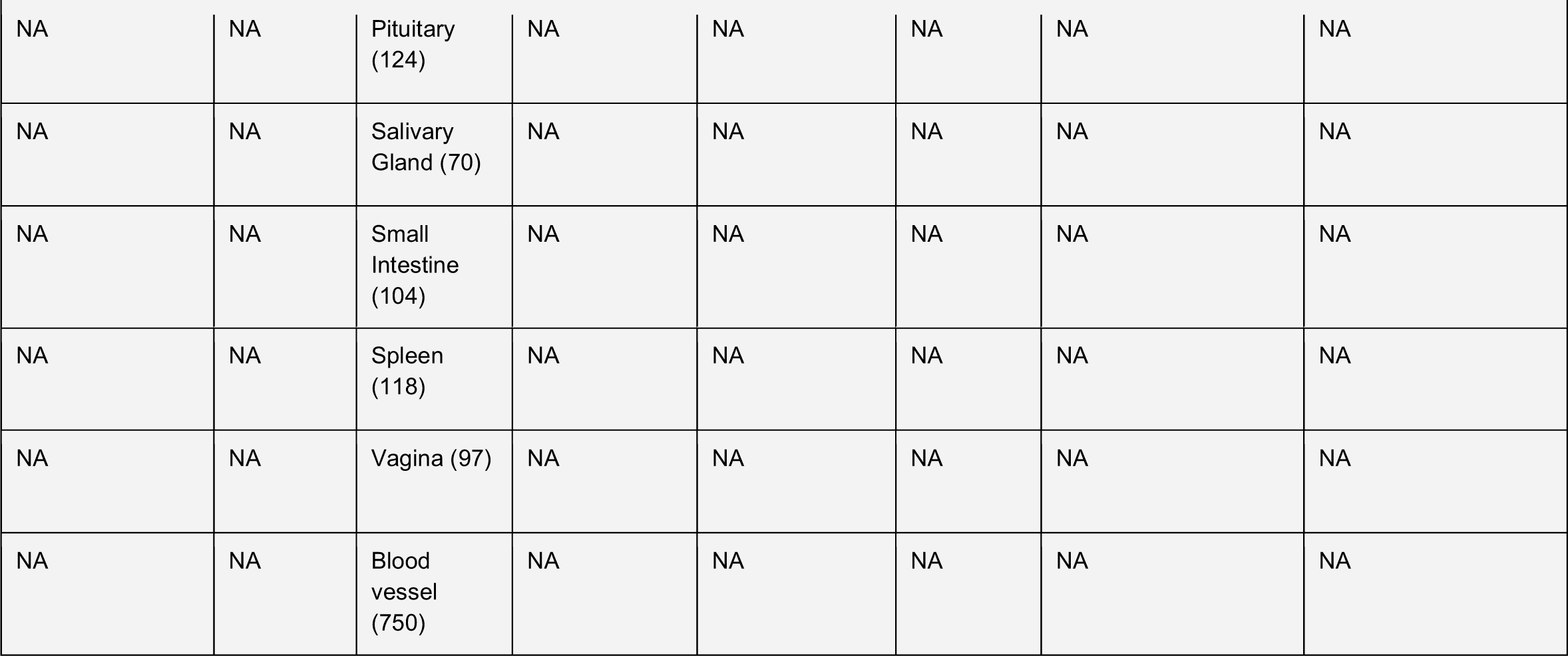
Sources and counts of tumor and tissue-matched normal samples.

**Table S2:** Percent of junctions not found in core normal samples, averaged across cancer types.

| SRA category: |  | Adult | Developmental | Stem Cells | Unexplained |
| --- | --- | --- | --- | --- | --- |
| Number of samples per SRA category required for set membership | 1 | 26.5% | 15.4% | 2.7% | 64.9% |
|  | 2 | 19.6% | 5.92% | 2.2% | 76.4% |

**Table S3:**
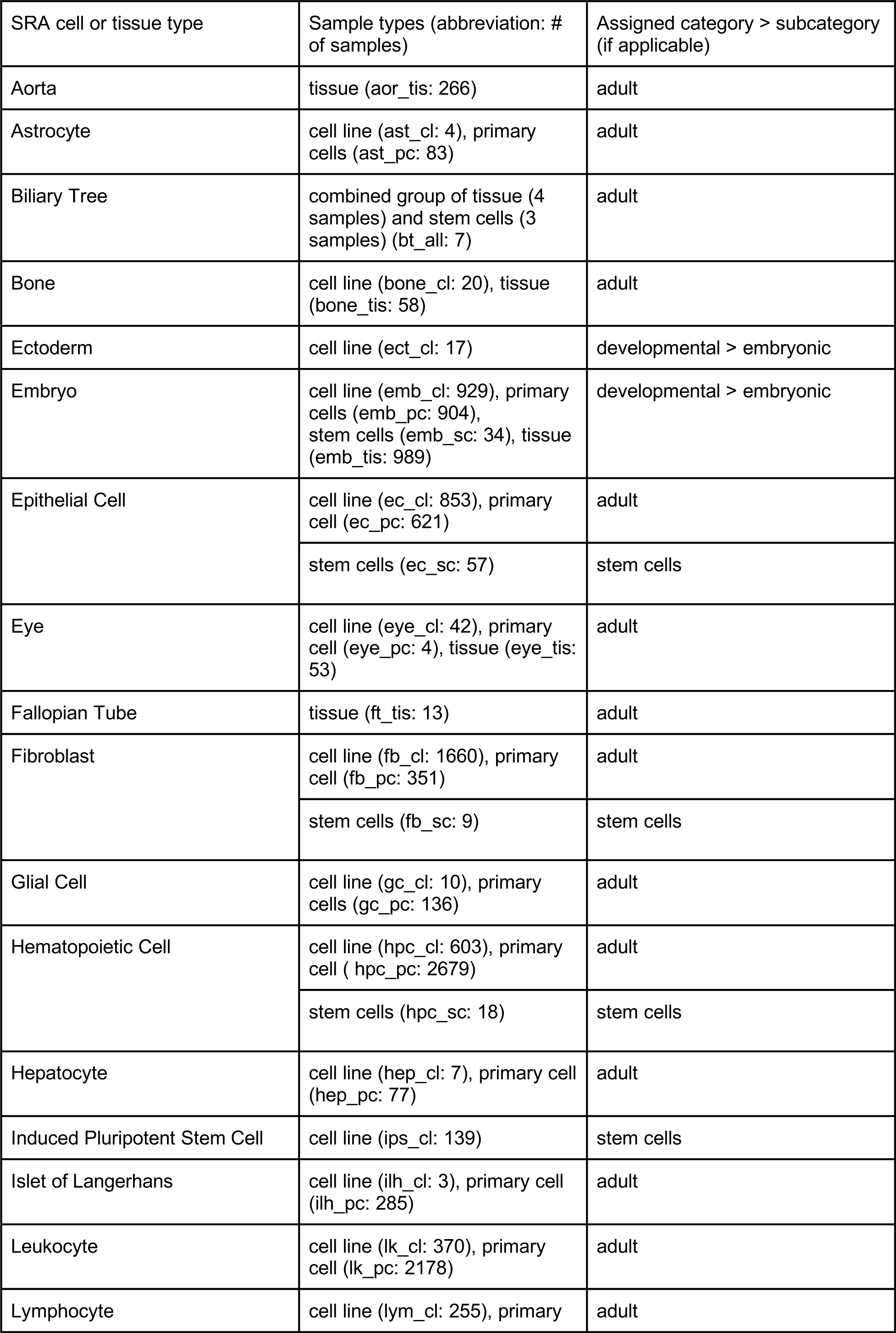
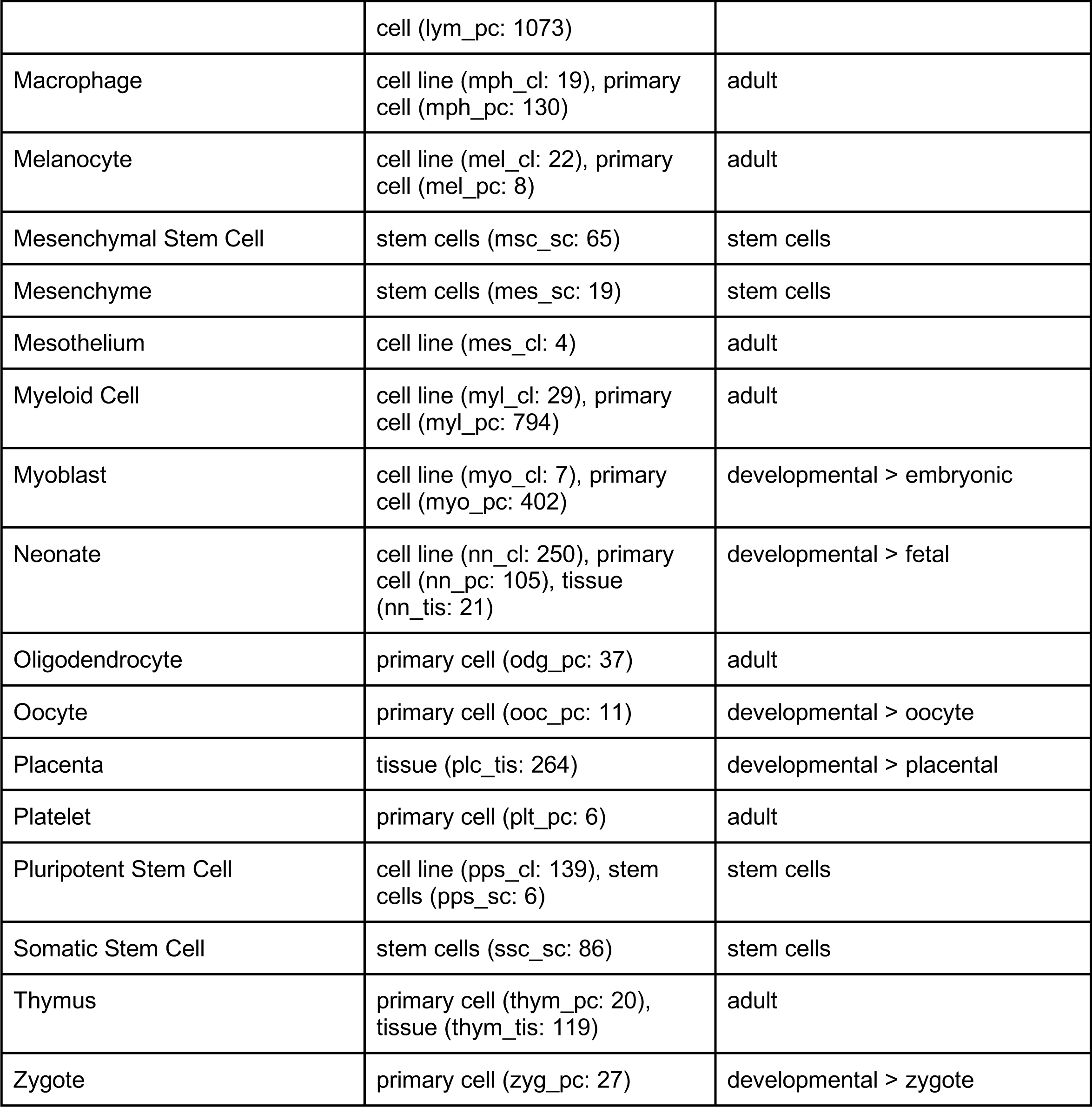
Selection of additional normal tissue and cell types analyzed.

**Table S4:**
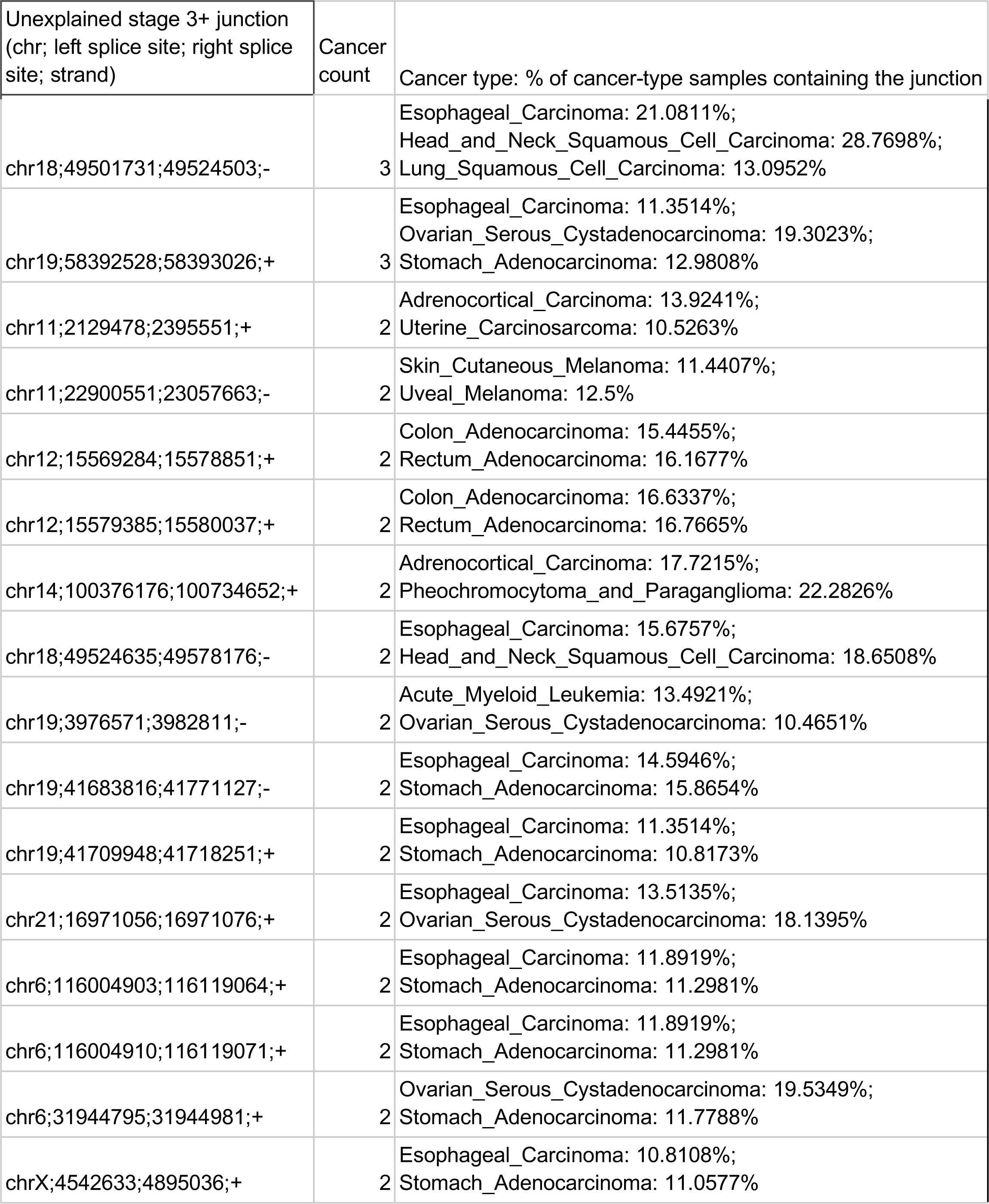
Unexplained junctions occurring in >10% of samples of multiple TCGA cancer types.

**Table S5:** Junction counts and proportion of antisense junctions for all TCGA cancer types

| TCGA cancer type | Core normals |  | Other adult non-cancer |  | Developmental |  | Stem cell |  | Unexplained |  |
| --- | --- | --- | --- | --- | --- | --- | --- | --- | --- | --- |
|  | junction count | antisense prevalence | junction count | antisense prevalence | junction count | antisense prevalence | junction count | antisense prevalence | junction count | antisense ratio |
| Acute Myeloid Leukemia | 1,745,361 | 27% | 183,003 | 46% | 46,261 | 48% | 4,580 | 45% | 427,768 | 47% |
| Adrenocortical Carcinoma | 941,404 | 20% | 7,830 | 39% | 2,369 | 38% | 326 | 33% | 22,053 | 44% |
| Bladder Urothelial Carcinoma | 2,401,749 | 24% | 58,874 | 35% | 17,652 | 31% | 3,348 | 26% | 155,124 | 42% |
| Brain Lower Grade Glioma | 2,847,326 | 26% | 64,093 | 40% | 19,716 | 38% | 3,129 | 32% | 165,091 | 41% |
| Breast Invasive Carcinoma | 4,013,058 | 27% | 170,106 | 36% | 44,855 | 36% | 7,520 | 30% | 441,104 | 37% |
| Cervical Squamous Cell Carcinoma and Endocervical Adenocarcinoma | 2,148,683 | 24% | 45,625 | 36% | 12,353 | 35% | 2,361 | 27% | 111,091 | 42% |
| Cholangiocarcinoma | 754,166 | 15% | 3,790 | 33% | 900 | 33% | 168 | 24% | 8,399 | 37% |
| Colon Adenocarcinoma | 2,228,830 | 25% | 64,988 | 37% | 19,894 | 31% | 2,841 | 32% | 155,589 | 42% |
| Esophageal Carcinoma | 3,939,154 | 29% | 349,351 | 43% | 113,466 | 45% | 13,813 | 38% | 1,039,014 | 46% |
| Glioblastoma Multiforme | 2,368,563 | 25% | 35,154 | 36% | 12,209 | 35% | 2,024 | 26% | 75,111 | 37% |
| Head and Neck Squamous Cell Carcinoma | 2,571,867 | 24% | 72,107 | 35% | 19,318 | 34% | 4,210 | 27% | 177,027 | 40% |
| Kidney Chromophobe | 1,042,201 | 22% | 10,778 | 42% | 3,081 | 37% | 490 | 26% | 28,961 | 42% |
| Kidney Renal Clear Cell Carcinoma | 2,925,458 | 26% | 93,435 | 38% | 21,535 | 38% | 3,160 | 32% | 197,822 | 40% |
| Kidney Renal Papillary Cell Carcinoma | 1,896,671 | 25% | 33,239 | 39% | 8,380 | 38% | 1,255 | 33% | 85,284 | 41% |
| Liver Hepatocellular Carcinoma | 1,889,979 | 23% | 43,871 | 36% | 9,302 | 33% | 1,486 | 27% | 91,086 | 40% |
| Lung Adenocarcinoma | 2,890,617 | 26% | 95,291 | 37% | 23,848 | 35% | 4,387 | 30% | 229,442 | 40% |
| Lung Squamous Cell Carcinoma | 3,018,232 | 24% | 92,301 | 33% | 27,381 | 30% | 5,604 | 22% | 212,501 | 36% |
| Lymphoid Neoplasm Diffuse Large B cell Lymphoma | 793,306 | 18% | 9,851 | 30% | 1,806 | 32% | 277 | 30% | 16,201 | 36% |
| Mesothelioma | 1,150,647 | 20% | 11,387 | 38% | 3,060 | 39% | 594 | 29% | 27,214 | 43% |
| Ovarian Serous Cystadenocarcinoma | 5,092,658 | 32% | 542,738 | 46% | 242,074 | 48% | 24,032 | 41% | 2,739,236 | 49% |
| Pancreatic Adenocarcinoma | 1,593,779 | 22% | 20,854 | 39% | 5,087 | 41% | 876 | 33% | 49,429 | 44% |
| Pheochromocytoma and Paraganglioma | 1,480,083 | 23% | 19,085 | 40% | 6,003 | 40% | 832 | 35% | 53,453 | 43% |
| Prostate Adenocarcinoma | 2,423,109 | 26% | 55,616 | 40% | 15,372 | 40% | 2,257 | 35% | 159,220 | 43% |
| Rectum Adenocarcinoma | 1,381,622 | 22% | 21,823 | 37% | 6,693 | 33% | 945 | 28% | 42,408 | 42% |
| Sarcoma | 1,985,824 | 23% | 31,671 | 35% | 10,306 | 33% | 1,699 | 28% | 77,762 | 39% |
| Skin Cutaneous Melanoma | 2,493,051 | 24% | 62,444 | 36% | 17,376 | 34% | 2,973 | 27% | 148,206 | 38% |
| Stomach Adenocarcinoma | 4,844,395 | 31% | 494,846 | 44% | 172,922 | 46% | 19,099 | 40% | 1,787,745 | 47% |
| Testicular Germ Cell Tumors | 1,627,329 | 19% | 26,027 | 26% | 12,435 | 23% | 5,054 | 17% | 49,836 | 31% |
| Thymoma | 1,364,454 | 21% | 19,938 | 32% | 4,892 | 31% | 1,057 | 26% | 40,450 | 37% |
| Thyroid Carcinoma | 2,501,295 | 27% | 54,285 | 42% | 14,124 | 42% | 2,217 | 37% | 147,977 | 44% |
| Uterine Carcinosarcoma | 993,014 | 18% | 8,062 | 35% | 3,328 | 30% | 601 | 29% | 22,474 | 39% |
| Uterine Corpus Endometrial Carcinoma | 2,499,341 | 26% | 70,746 | 36% | 20,596 | 35% | 3,226 | 31% | 155,889 | 41% |
| Uveal Melanoma | 849,072 | 20% | 7,281 | 40% | 1,799 | 42% | 276 | 35% | 16,284 | 42% |

